# CRISPR-mediated activation of biosynthetic gene clusters for bioactive molecule discovery in filamentous fungi

**DOI:** 10.1101/2020.01.12.903286

**Authors:** Indra Roux, Clara Woodcraft, Jinyu Hu, Rebecca Wolters, Cameron L.M. Gilchrist, Yit-Heng Chooi

## Abstract

Accessing the full biosynthetic potential encoded in the genomes of fungi is limited by the low expression of most biosynthetic gene clusters (BGCs) under common laboratory culture conditions. CRISPR-mediated transcriptional activation (CRISPRa) of fungal BGC could accelerate genomics-driven bioactive secondary metabolite discovery. In this work, we established the first CRISPRa system for filamentous fungi. First, we constructed a CRISPR/dLbCas12a-VPR-based system and demonstrated the activation of a fluorescent reporter in *Aspergillus nidulans*. Then, we targeted the native nonribosomal peptide synthetase-like (NRPS-like) gene *micA* in both chromosomal and episomal contexts, achieving increased production of the compound microperfuranone. Finally, multi-gene CRISPRa led to the discovery of the *mic* cluster product as dehydromicroperfuranone. Additionally, we demonstrated the utility of the variant dLbCas12a^D156R^-VPR for CRISPRa at room temperature culture conditions. Different aspects that influence the efficiency of CRISPRa in fungi were investigated, providing a framework for the further development of fungal artificial transcription factors based on CRISPR/Cas.

---

Fungal genome mining has emerged as a promising strategy for the discovery of novel bioactive secondary metabolites (SMs) (also known as natural products).^1,2^ Genomic surveys have revealed that fungal species typically harbor 30–80 biosynthetic gene clusters (BGCs) each encoding the biosynthetic pathway required to produce a SM(s).^3^ However, the vast majority of BGCs remain uncharacterized or ‘cryptic’ as the products they encode are undetectable under standard culture conditions, often because BGCs remain ‘silent’ or lowly expressed due to tight regulatory control.^1,4^ Filamentous fungi, which have yielded a plethora of SMs with pharmaceutical and agricultural applications^5^, thus serve as attractive targets for genome mining of novel molecules.

Improved understanding of SM biosynthesis has led to the development of various bioinformatic tools and strategies for the prioritization of BGCs for genome mining, increasing the chance of discovery of novel molecules or molecules with desired bioactivities.^6,7^ Current strategies for activating specific BGCs typically involve promoter exchange of individual genes in the BGC with strong promoters and marker recycling for sequential chromosomal manipulation.^8^ About half of the BGCs in fungi contain a gene encoding a pathway-specific transcription factor (TF)^1^, in which activation of BGC expression could be achieved by overexpression of the TF.^9^ However, in many cases TF overexpression does not result in successful BGC activation, which has prompted strategies such as engineering of hybrid TFs.^10^ On the other hand, reconstruction of the pathway in a heterologous host is the preferred method for BGCs from fungi that are genetically intractable.^5^ Filamentous fungi are the most compatible heterologous hosts for expression of fungal BGCs, not requiring intron removal or codon optimization.^11^ For example, *Aspergillus nidulans* has been successfully utilized as a heterologous host by several groups,^12–14^ including ours.^15,16^ However, reconstruction of BGCs require the cloning of multiple and often large biosynthetic genes,^16^ which can be cumbersome. Other methods to express BGCs as engineered polycistronic mRNA also rely on amplification or synthesis of all BGC genes.^17^

To access cryptic SMs more efficiently and pave the way to higher-throughput pathway-specific genome mining for drug discovery, new tools for transcriptional activation are necessary. Inspired by pathway-specific TFs, we aim to develop a CRISPR-mediated transcriptional activation (CRISPRa) approach for accessing the BGCs in filamentous fungi (Figure 1a). Unlike CRISPR genome editing that uses nuclease-active CRISPR/Cas for targeted DNA cleavage, in CRISPRa the CRISPR/Cas complex is nuclease-deactivated and repurposed as an artificial transcription factor to modulate gene expression. CRISPRa systems make use of activation effectors linked to a nuclease deactivated Cas protein (dCas) in complex with a short guide RNA containing a programmable ~20 nt sequence complementary to a target site within a gene regulatory region to achieve transcriptional activation. ^18,19^ The activation of multiple genes can be achieved with multiplexed CRISPRa, in which multiple guide RNAs are expressed simultaneously.^20^ CRISPRa has already been used to tune the expression of biosynthetic pathway coding genes,^21,22^ including in ascomycetous yeasts. ^23,24^ Although CRISPR/Cas genome editing has been widely adopted in filamentous fungi, ^25–28^ to our knowledge, CRISPRa has not been demonstrated in these organisms and has not been used as a tool for compound discovery.

**Figure 1.**
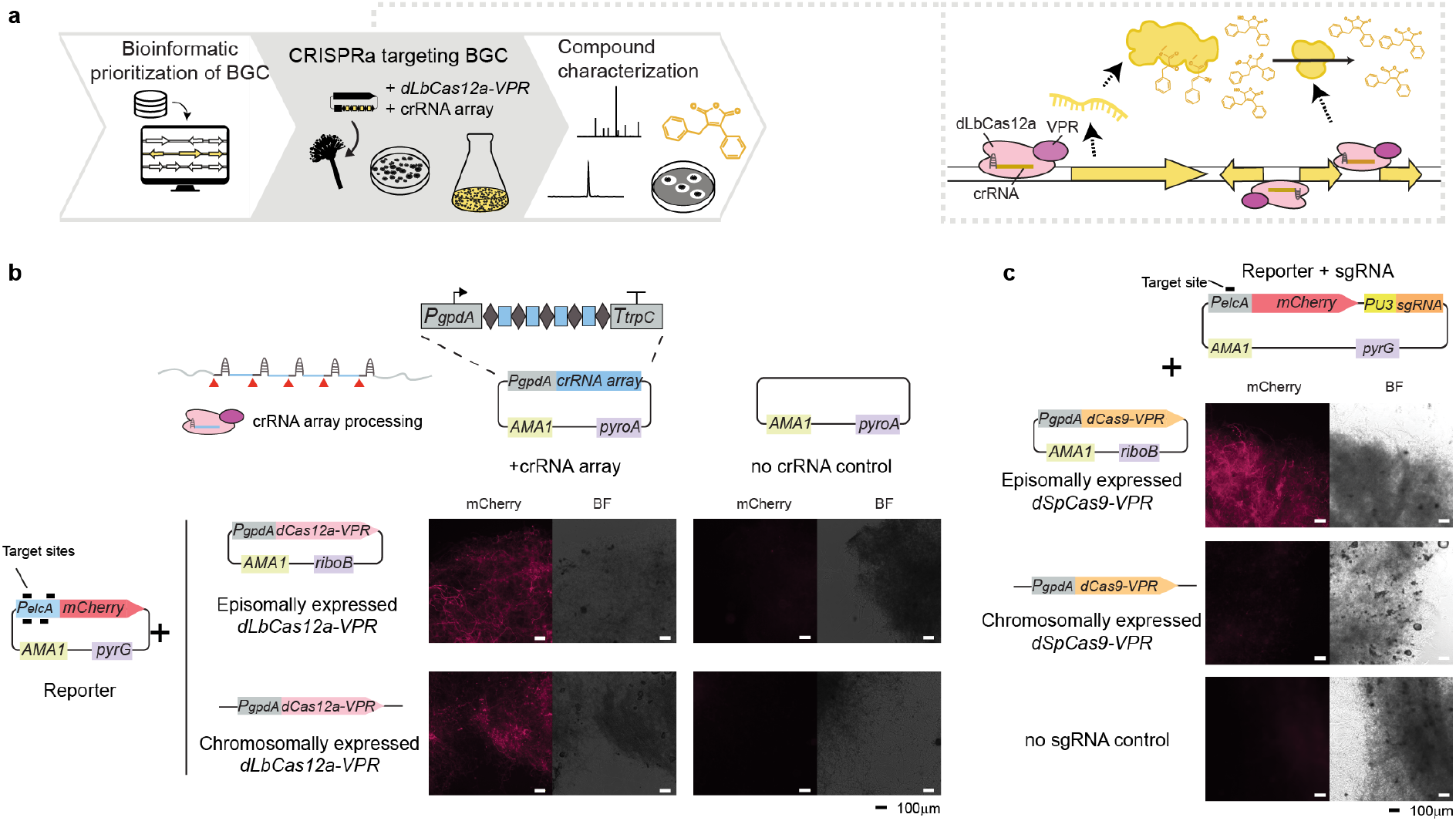
Proof-of-concept for fungal CRISPRa **a.** Schematic of a CRISPRa-based genome mining pipeline. After BGC bioinformatic prioritization, the designed crRNAs are rapidly assembled by Type IIs cloning from annealed oligos in an expression vector and transformed into the fungal host along with the dCas effector. The CRISPRa complex is targeted to the regulatory regions of the selected BGC, upregulating gene expression and consequently increasing production of the encoded compound(s). Higher titers facilitate compound detection, screening for bioactivity and further chemical characterization. **b.** CRISPR/dLbCas12a-VPR mediated activation of *P_elcA_-mCherry* in *A. nidulans.* Representative fluorescent microscopy images of CRISPRa transformant mycelia demonstrate consistent *mCherry* reporter activation, which implies the processing of crRNA array from a *P_gpdA_*-derived RNAPII-driven transcript by dLbCas12a-VPR. Activation is observed both episomally and chromosomally encoded dLbCas12a-VPR systems (Figure S1). **c.** Activation strength of the CRISPR/dSpCas9-VPR system showed dependency on dSpCas9-VPR expression strategy (Figure S2). In all microscopy images mycelia were observed under brightfield (BF) and mCherry filter after overnight growth on stationary liquid culture at 37 °C. Scale bar 100 μm.

In this work, we developed a suite of fungal CRISPRa vectors and test them in *A. nidulans*. We used dCas proteins fused to the synthetic tripartite activator VP64-p65-Rta (VPR), ^19^ which contains a fusion of three activation domains and has led strong activation in several species,^29^ but untested in filamentous fungi. We built and tested two different CRISPRa systems, based on dLbCas12a from *Lachnospiraceae bacterium* (previously known as LbCpf1)^18^ and dSpCas9 from *Streptococcus pyogenes.*^19^ SpCas9 is currently the more widely used system in fungal CRISPR applications, but the delivery of multiple single guide RNAs (named sgRNAs) requires several promoters or additional engineering for RNA processing. ^20,28^ The alternative system driven by LbCas12a has the potential to simplify multiplexing due to its use of a shorter guiding CRISPR RNA (named crRNA) and the capability of Cas12a to process multiple crRNAs from a single precursor crRNA array.^30^ By targeting a BGC (*mic* cluster encoding a nonribosomal peptide synthetase-like enzyme) in *A. nidulans*, we demonstrated the potential of CRISPR/dLbCas12a-VPR as a tool for fungal BCG activation and accelerating compound discovery. We evaluated aspects that influence CRISPR/dLbCas12a-VPR efficiency for biosynthetic gene activation, such as CRISPRa component delivery, targeting parameters and culture temperature, and explore alternatives to overcome some of the limitations discovered.

## Results and Discussion

### Construction and testing of fungal CRISPRa systems

To develop a CRISPRa system for filamentous fungi, we constructed and tested CRISPR/dLbCas12a-VPR- and CRISPR/dSpCas9-VPR-based systems in the model organism and chassis *A. nidulans.* To evaluate alternative strategies for expressing either dCas effector, we created parent strains with a chromosomally integrated *dCas-VPR* expression cassette and compared their performance with entirely AMA1-episomally encoded systems. The AMA1 sequence acts as an extrachromosomal vector replicator and confers increased transformation frequency in several filamentous fungi species^31^. AMA1-bearing vectors are found at multiple copies per nucleus although their genetic stability has been reported to be limited under non-selective conditions. We built on the triple auxotrophic mutant *A. nidulans* LO8030,^32^ which can maintain AMA1 vectors by complementation with the selectable markers *pyrG*, *riboB*, *pyroA*.^16^ The AMA1 vector set allowed us to separate the key components of the CRISPRa system into different vectors for modularity during the initial validation. As a proof-of-concept target, we built a fluorescent reporter by fusing *mCherry* to *Parastagonospora nodorum elcA* promoter (*P_elcA_*), which belongs to a ‘silent’ polyketide synthase gene, ^33^ and delivered it encoded on a AMA1 vector.

Firstly, we assessed dLbCas12a-VPR crRNA array processing capability in *A. nidulans*. As individual crRNAs can be excised by LbCas12a from arrays embedded in RNA polymerase II (RNAPII)-driven transcripts^34^ we built a crRNA array expression cassette with *gpdA* promoter (*P_gpdA_*) and *trpC* terminator (*T_trpC_*) (Note S1), which are parts widely portable across fungal species.^35^ Due to the lack of characterization of the transcription start site (TSS) of the *elcA* gene, we designed four crRNAs targeting a window 88–327 bp upstream of the open reading frame start codon (Figure 1b). We delivered the four-crRNA array on an AMA1-pyroA vector co-transformed with the *P_elcA_* reporter. After growing mycelial mass, we observed activation of *mCherry* expression in the CRISPRa transformants compared to the no crRNA control, in both chromosomally and episomally expressed dLbCas12a-VPR systems (Figure 1b, Figure S1). The results demonstrated the viability of the RNAPII-promoter *P_gpdA_* to deliver LbCas12a crRNA arrays.

In parallel, we built and tested a dSpCas9-VPR system, with a sgRNA expression cassette driven by the RNA polymerase III (RNAPIII) promoter U3 from *Aspergillus fumigatus* (*AfP_U3_*)^28^ (Note S1). In this case, four sgRNA were tested individually, targeting a window 162–342 bp from the reporter start codon, and delivered in a single AMA1-pyrG vector together with the reporter construct *P_elcA_-mCherry*. We observed that the system with chromosomal expression of *dSpCas9-VPR* resulted in activation levels only noticeable at prolonged exposure times, while the system with episomally expressed dSpCas9-VPR resulted in stronger fluorescence (Figure 1c, Figure S2a–b). A possible interpretation is that the single-copy chromosomal *dSpCas9-VPR* cassette failed to achieve expression levels above the required threshold for strong observable activity, making the multicopy AMA1-encoded system more effective in comparison. We also observed that the activation level was stronger for the sgRNAs targeting closer to the target gene start codon (Figure S2b). CRISPR-mediated activation has been reported to be stronger targeting closer to the upstream region of a gene TSS,^36^ but in this case, we lack information about *elcA* TSS. We further attempted to deliver sgRNAs from an independent AMA1-pyroA vector, but when co-transformed with the reporter vector and the dSpCas9-VPR expression vector, fluorescence was not observed (Figure S2c).

When comparing dCas9- and dCas12a-driven systems, CRISPR/dLbCas12a-VPR presented advantages in multiplexing capability and supported expression of CRISPRa components in various configurations. Additionally, we showed that dCas12a can process crRNA transcripts driven by the commonly used RNAPII promoter P_*gpdA*_. This increased the potential portability of the CRISPRa system across many fungal species and opens up the possibility of driving crRNA expression under a variety of characterized RNAPII promoters, including inducible promoters (eg. alcohol-inducible P_*alcA*_). On the other hand, multiplexing with Cas9 sgRNAs would require expression from multiple cassettes driven by RNAPIII promoters or additional RNA processing mechanisms if expressed as an array driven by RNAPII promoters.^20^ For these reasons, we further explored the activation of biosynthetic genes in *A. nidulans* using the CRISPR/dLbCas12a-based CRISPRa system.

### CRISPR/dLbCas12a-VPR mediated activation of *micA* synthetase gene increases microperfuranone production

To test whether CRISPR/dLbCas12a-VPR mediated activation of a fungal biosynthetic gene could induce metabolite production, we targeted the native *A. nidulans micA* (*AN3396*) gene which encodes a nonribosomal peptide synthetase-like (NRPS-like) enzyme responsible for the biosynthesis of microperfuranone (**1**) (Figure 2a). The biosynthetic function of *micA* had been previously decoded by using promoter replacement strategy,^37^ after unsuccessful attempts to elicit the biosynthesis of cryptic NRPS-like products by varying *A. nidulans* culture conditions.

**Figure 2.**
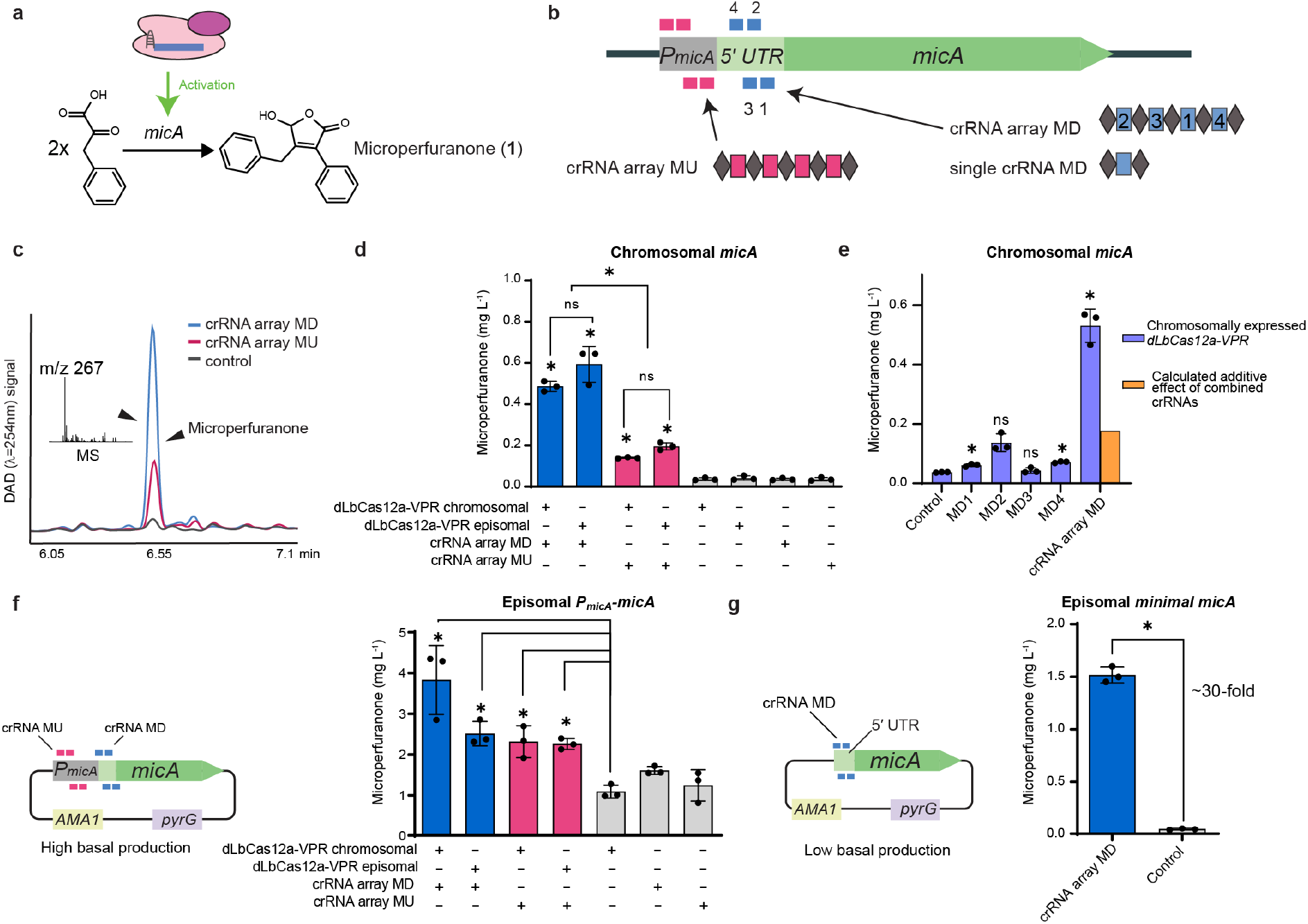
CRISPRa-mediated microperfuranone (**1**) production. **a.** Upregulation of *micA* results in the biosynthesis of **1** from two phenylpyruvate molecules ^37^. **b.** Chromosomal *micA* gene with individual crRNA binding sites shown in magenta (crRNA array MU) and blue (crRNA array MD). Numbers assigned to MD crRNAs are indicative of targeting position in respect to *micA* start codon, with their position in the crRNA array MD also indicated in the scheme. **c.** DAD (λ=254 nm) chromatograms of *A. nidulans* culture media extracts show increases in the peak identified as **1** in strains with CRISPR-mediated activation of *micA* (magenta and blue) as compared to the control with no crRNA (grey). Expected mass of **1** is observed as main ion in the peak by MS. **d.** CRISPR-mediated activation of chromosomal *micA*. All CRISPRa strains showed a significant increase in the production of **1** compared to their respective dLbCas12a-VPR control with no crRNA. Targeting CRISPRa with MD crRNA array (blue) resulted in significantly higher production of **1** compared to targeting with MU crRNA (magenta). There was no significant difference between both dLbCas12a-VPR expression strategies. **e.** When targeting *micA* with single MD crRNAs low or no activation is observed compared to the no crRNA control. The low calculated additive of single crRNA CRISPRa (described in Methods) suggests a synergistic activation effect when making use of the four-crRNA array MD. **f.** Increases in the titer of **1** are observed in media extracts of CRISPRa strains with extra episomal *micA* copies (high basal production) compared to the no crRNA control in strains with episomal *micA* vector. **g.** In strains harboring episomal copies of a shorter *micA* variant (low basal production), CRISPRa increased the production of **1** by ~30-fold compared to the control. In all the figures, calculated titer (mg L^−1^) values are the mean of three biological replicates which specific values are indicated as black dots, error bars represent SD. Two-sided Welch’s T-test with Holm-Šídák multiplicity correction per figure was performed. Asterisk indicates corrected P-value<0.05, (ns) not significant. Individual P-values are listed in Table S9.

We first targeted *micA* with multiple crRNAs to increase the likelihood of achieving strong activation, an approach used in CRISPRa screenings.^38^ We followed previously devised guidelines for CRISPRa in eukaryotes,^36^ and targeted a region 119–303 bp upstream of *micA* TSS with a four-crRNA array named MU (*micA* Upstream TSS) (Figure 2b). To explore the utility of CRISPRa for genes that lack TSS information, which is the case for 59% of *A. nidulans* BGC genes (Figure S3a), we also tested an alternative TSS annotation-blind targeting criteria, taking the gene start codon as reference. Given that most *A. nidulans* BGC genes have a short 5′ untranslated region (UTR) (Figure S3b), we targeted a window 139–324 bp upstream of the *micA* start codon with a four-crRNA named MD (*micA* Downstream TSS), which in the case of *micA* corresponds to the 5′ UTR. Analysis by liquid chromatography coupled to a diode array detector and mass spectrometer (LC-DAD-MS) showed increases in the production of **1** in media extracts from all CRISPRa transformants when compared to the background levels in the controls (Figure 2c). Interestingly, targeting the 5′ UTR of *micA* with crRNA array MD resulted in significantly higher production than when targeting the *micA* promoter region with crRNA array MU, reaching titers of **1** up to 0.6 mg L^−1^ (Figure 2d). Nevertheless, targeting with MU still led to a ~4.5-fold increase in production with a titer up to 0.2 mg L^−1^ compared to the controls. No significant difference in performance was observed when comparing between chromosomally and episomally expressed dLbCas12a-VPR systems for both crRNA arrays.

In order to enable rapid cloning of different crRNAs for further testing, we established a domesticated version of the AMA1-pyroA expression vector, which allowed one-step Type IIS cloning of crRNA arrays using annealed oligonucleotides (Figure S4). We verified the null effect of P_*gpdA*_ domestication (Figure S4b).

To examine the effect of each crRNA in *micA* activation, individual crRNAs from MD and MU arrays were delivered in strains harboring chromosomally integrated *dLbCas12a-VPR* (Figure 2e, Figure S5). We observed a minimal increase in the production of **1** when targeting with some MD crRNA, while in most cases the production of **1** was indistinguishable from the no crRNA control. The notably difference between the production of **1** in strains with single crRNAs and with multiple-crRNA array CRISPRa is indicative of a synergistic activation effect (Figure 2e). Thus, in this case, the synergistic multi-crRNA activation of *micA* is required for achieving metabolite production of **1**.

To test whether the production of **1** could be further increased, we targeted *micA* with the crRNAs from both MD and MU crRNA arrays simultaneously. To this end, we re-cloned the crRNA array MD into an AMA1-pyrG vector to allow co-transformation with the crRNA array MU encoded on an AMA1-pyroA vector. Co-transformation of MD and MU crRNA arrays for *micA* activation resulted in further increase in the production of **1** (up to ~0.8 mg L^−1^) (Figure S6). Interestingly, the crRNA array MD alone delivered from the AMA1-pyrG vector resulted in a considerable increase in the titer of **1** compared to when delivered using AMA1-pyroA vector, which might contribute to the dual crRNA array increased production (Figure S6).

Finally, to evaluate the broader utility of CRISPRa targeting episomal genes in *A. nidulans*, we co-transformed additional copies of *micA* encoded on an AMA1 vector. The transformants harboring episomal copies of *micA* with its full-length promoter showed relatively high basal production of **1** even in the absence of CRISPRa (Figure 2f). However, we still observed a consistent increase in the titers of **1** when CRISPRa of *micA* was performed with either MD or MU crRNA arrays, reaching up to over 4 mg L^−1^ (Figure 2f). We further tested targeting a shorter episomal *micA* variant with low basal production of **1**. When co-transforming with MD, targeting the still-present 5′ UTR, we observed the largest activation fold change with a ~30-fold increase in the production of **1** compared to the control, reaching final titers of ~1.5 mg L^− 1^ (Figure 2g). Taken together, these results show that CRISPRa can affect the expression of episomally encoded genes, as the increase in the production of **1** is not explained by the activation of chromosomal *micA* alone.

Considering filamentous fungi genomes in public databases often lack 5′ UTR annotations, the viability of selecting crRNA targets in a TSS annotation-blind manner is a significant advantage. In the case of *micA*, targeting a short distance upstream of the gene start codon, despite falling in the 5′ UTR, resulted in successful production of microperfuranone. This is surprising, given that the binding of a CRISPRa complex downstream of the TSS has been proposed to act as a roadblock of transcription,^23^ however our observations indicate that the binding of dLbCas12a-VPRs to the *micA* 5′ UTR region redefines the local transcriptional landscape by other means with the resulting activation of this locus. ^39^ Although this may be a locus-specific effect, our results suggest that a TSS annotation-blind targeting criteria could be a viable alternative when targeting genes with incomplete gene annotation, as demonstrated for *micA* and also above, for *P_elcA_* driving *mCherry* expression.

### Multi-gene activation uncovered cryptic biosynthetic gene cluster product

The gene *micA* (*AN3396*) has been proposed to belong to a multigene BGC^37,40,41^, in which the final product has remained uncharacterized. Besides *micA*, the *mic* cluster consists of a putative cytochrome P450 (*AN3394*) and a hypothetical gene (*AN3395*), referred to as *micB* and *micC*, respectively and lacks a pathway-specific TF.

To assess the feasibility of performing simultaneous activation of multiple genes with CRISPR/dLbCas12a-VPR, we aimed to co-activate the proposed *mic* cluster. We co-transformed the four-crRNA array MD targeting *micA* along with a second crRNA array targeting *micB* and *micC* (Figure 3a). This new three-crRNA array, named crRNA array P, targeted a window of 209bp in the middle of the short 396 bp bi-directional promoter between the divergently oriented *micB* and *micC* genes. We delivered the two crRNA arrays from independent AMA1 episomal vectors in strains harboring chromosomally integrated *dLbCas12a-VPR* (Figure 3a). To account for the above-mentioned influence of the crRNA delivery vector selection marker on CRISPRa strength, we tested both dual vector delivery combinations with *pyrG* and *pyroA* selection markers (Figure 3a).

**Figure 3.**
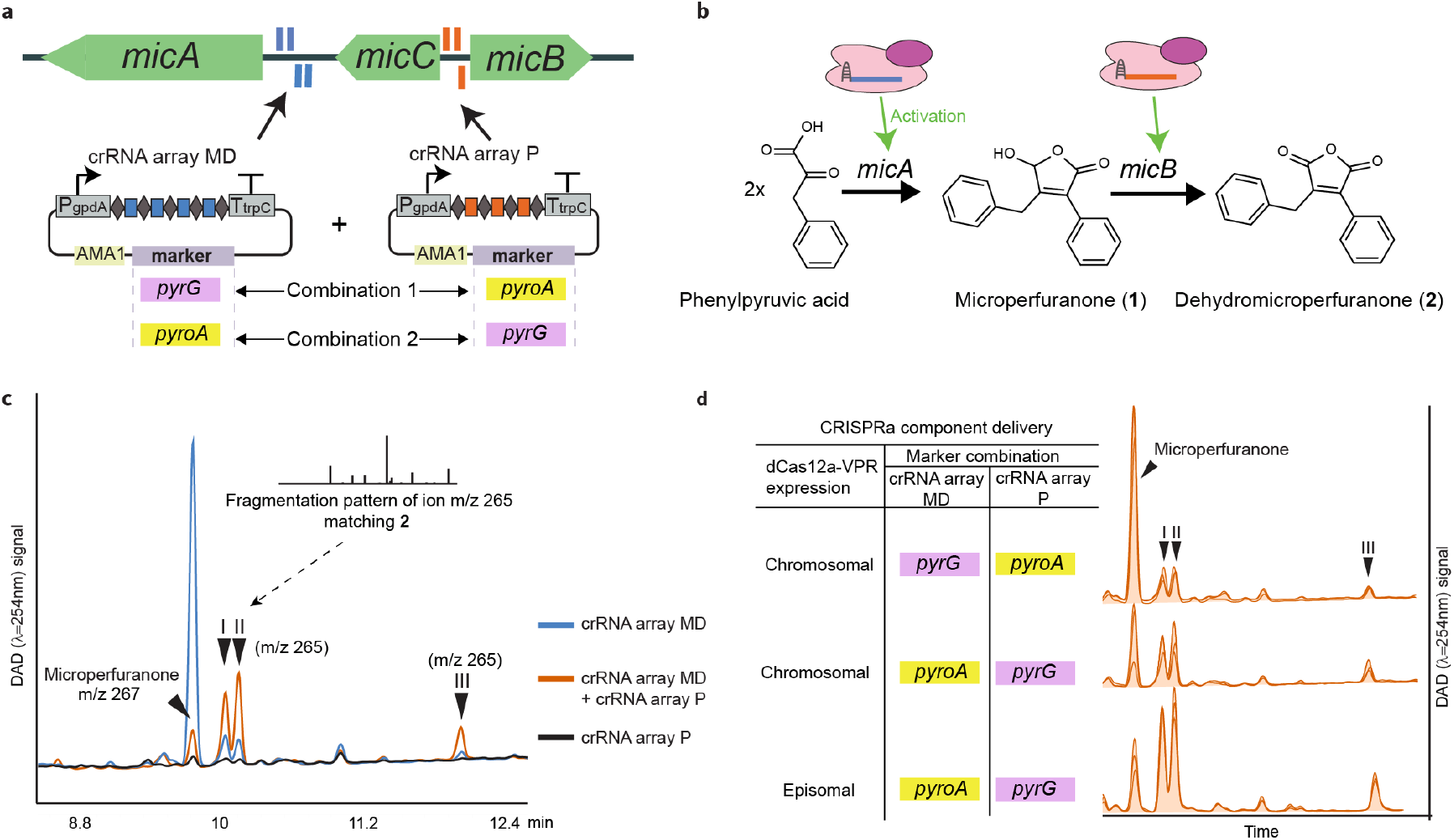
Elucidating the *mic* cluster final product with multi-gene activation **a.** Schematic of the experimental set-up of *mic* cluster activation with two-vector crRNA arrays delivery and different markers combinations of *pyrG* (purple) and *pyroA* (yellow). The *mic* cluster genes are shown in green alongside the target sites of crRNA MD array (blue) and crRNA P array (orange). **b.** Proposed dehydromicroperfuranone (**2**) structure and biosynthetic pathway. **c.** Representative overlaid DAD (λ=254 nm) chromatograms of media extracts from strains with both crRNA arrays MD and P (orange), crRNA array MD (blue) and crRNA array P (black). Multi-gene CRISPRa results in the increase of the peaks I–III whose main ion *m*/*z* 265 yielded a fragmentation pattern matching **2** by LC-MS/MS analysis (Table S1). **d**. Production of each peak in multiple activation strains is dependent on CRISPRa component delivery strategy. The different DAD (λ=254 nm) chromatograms represent three biological replicates per delivery strategy (Figure S7). We observe that the production of the peaks I–III is favored in the marker combination crRNA MD in an AMA1-pyroA vector and crRNA P in an AMA1-pyrG vector.

LC-DAD-MS analysis of *A. nidulans* culture extracts showed that both multiplexed-CRISPRa strains presented a decrease in the precursor microperfuranone (**1**) and an increase in the size of three new peaks detected by DAD and MS, arbitrarily named peaks I, II and III (Figure 3c, Figure S7a–b). We also observed that the magnitude of the changes in the metabolic profile were dependent on the marker combination used in the delivery of the crRNA arrays. The delivery of crRNA P on an AMA1-pyrG vector and crRNA MD in an AMA1-pyroA vector favored the production of the peaks I-III (Figure 3d). We further tested this crRNA array delivery combination switching to episomally encoded *dLbCas12a-VPR* and observed an increased production of the peaks I–III relative to the peak from **1** (Figure 3d, Figure S7c). Taken together these results indicate that multiple gene CRISPRa can be used to explore the metabolite products of a cryptic BGC and that the activity can be tuned to favor the final product of the cluster.

The observed mass of the ions accumulated in the peaks I-III was m/z 265 [M+H]^+^, 2 Da less than the molecular mass of **1**, suggesting that an oxidation had occurred. Searching the chemical literature for structures related to **1** corresponding to a mass of 264 Da led us to a previously reported metabolite, 3-carboxy-2,4-diphenyl-but-2-enoic anhydride, herein renamed as dehydromicroperfuranone (**2**) (Figure 3b), which was first isolated from *A. nidulans* IFO 6398 as a plant growth promoting compound.^42^ Indeed, we observed that the MS/MS fragmentation pattern of the 265 m/z ions shared almost all fragment masses with the predicted spectra for **2** by CFM-ID ^43^ (Table S1).

As the genetic basis for biosynthesis of **2** was not previously identified, to ensure that the production of the peaks I–III is due to CRISPRa co-targeting the *micB–C* promoter, we verified the final product of the *mic* cluster by promoter replacement of *micA*, *micB* and *micC*. When expressing *micA* and *micB* from an alcohol dehydrogenase inducible promoter (*PalcA*), the metabolic profile presented the peaks I–III as observed by CRISPRa, although the metabolites were produced in higher quantities (Figure S8a). The co-expression of *micA*–*C* resulted in the same metabolic profile as *micA*–*B* (Figure S8a). This revealed the function of MicB as a cytochrome P450 monooxygenase responsible for converting a secondary alcohol on **1** to a ketone group, forming a maleic acid anhydride moiety.

To corroborate the structure of the compounds, we attempted to purify the peaks I–III. Due to increased polarity, the peaks were only extractable from the culture medium with acidified ethyl acetate or adsorbent resin (Figure S8b), which might explain why it have not been observed in previous studies that detected *micB* expression but used ethyl acetate extraction for metabolite profiling of *A. nidulans*.^37,44^ The peaks I-II co-eluted during semi-preparative HPLC purification, while peak III could be isolated as a single peak. Surprisingly, the ^1^H-NMR and ^13^C-NMR spectra of the peaks I-II mixture and peak III in deuterated chloroform appear to be identical and matched the previously reported chemical shifts for **2** ^42^ (Table S2 and Figures S9–12). When reconstituting the NMR sample in methanol for analysis by LC-DAD-MS, the purified peaks reverted to multiple peaks (Figure S8c) and we further observed that the samples also existed as mixtures when analyzed by NMR in deuterated methanol (Figures S13–14). These results suggest that the compounds in the peaks I-III are interchangeable tautomeric/cis-trans isomers or opened/closed ring forms of **2** in acetonitrile or methanol (Figure S8d) but exist as a single entity in chloroform during NMR analysis (Table S2).

Taken together, the results from LC-MS/MS and NMR analysis support **2** as the metabolite product of the *mic* cluster. The results also conclusively demonstrated that multi-gene activation can be achieved by CRISPR/dLbCas12a-VPR. Here, we delivered each crRNA array from separate vectors for keeping the modularity in initial testing. However, longer crRNA arrays could be used to target a whole BGC, with encouraging precedents in the recent literature of Cas12a-mediated processing of 25 crRNAs from a single transcript and simultaneous upregulation of ten genes.^45^

### The variant dLbCas12a^D156R^ improves activation in cultures at 25 °C

Cas12a systems are known to exhibit reduced activity below 28°C,^46^ making its implementation troublesome in ectotherm animals and some plants. This could also compromise the wider applicability of fungal CRISPRa, as the majority of fungi have optimum growth temperatures between 25 °C and 30 °C,^47^ and the production of some SMs is favored at room temperature.^1^ In our laboratory, we routinely express heterologous biosynthetic genes from non-thermotolerant fungi in *A. nidulans* at 25 °C for metabolite production.^15,16^ To evaluate CRISPRa performance at temperatures lower than 37 °C, we investigated CRISPR/dSpCas9-VPR and CRISPR/dLbCas12a-VPR mediated activation of the *P_elcA_-mCherry* reporter across multiple temperatures. The fluorescence observed at 30 °C was comparable to the samples grown at 37 °C in both systems. However, at room temperature, low fluorescence was observed in CRISPR/dSpCas9-VPR samples and not detectable in CRISPR/dLbCas12a-VPR samples (Figure S15).

Due to the restrictions on dLbCas12a-VPR activity at room temperature (25 °C) a putative temperature tolerant variant was investigated. Taking inspiration from the AsCas12a^E174R^ variant, recently reported to possess increased double stranded DNA cleavage efficiency *in vitro* at 25 °C,^48^ we built an LbCas12a mutant harboring the homologous mutation D156R identified by aligning the AsCas12a/LbCas12a crystal structures^49,50^ (Figure S16).

We tested the variant dLbCas12a^D156R^-VPR targeting chromosomal *micA* with the crRNA array MD. We observed significant CRISPR/dLbCas12a^D156R^-VPR-mediated activation at 25 °C, a temperature at which CRISPR/dLbCas12a-mediated activation was not observed (Figure 4b). However, at 37 °C CRISPR/dLbCas12a^D156R^-VPR achieved lower final production of **1** than the original dLbCas12a-VPR system (Figure 4b). We also observed evidence of CRISPR/dLbCas12a^D156R^-VPR-mediated fluorescence activation at 25 °C (Figure 4a, Figure S15b). During the preparation of this manuscript, the LbCas12a^D156R^ variant was reported to exhibit increased low-temperature genome editing efficiency *in vivo* ^51^. Here, we demonstrated that the temperature tolerance property is translatable to CRISPRa.

**Figure 4.**
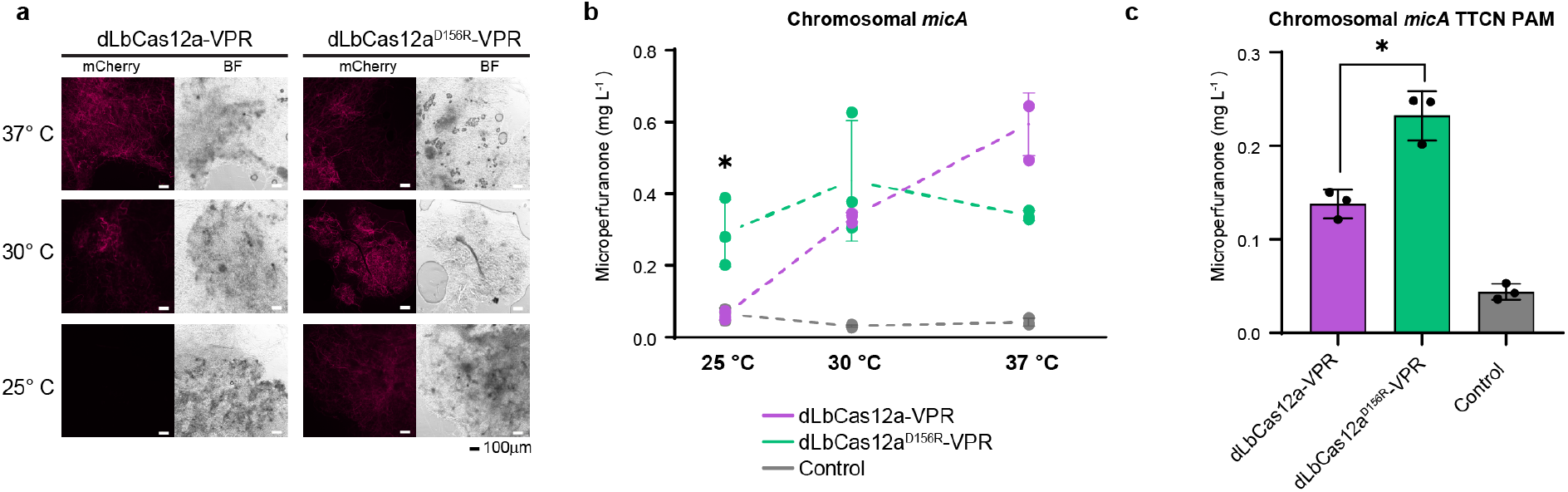
The variant dLbCas12a^D156R^-VPR outperforms at some limiting conditions for dLbCas12a-VPR. **a.** Representative microscopy images of *A. nidulans* mycelia grown at different temperatures show that CRISPR/dLbCas12a-VPR mediated activation of the fluorescent reporter *P_elcA_-mCherry* is restricted at 25 °C. The variant dLbCas12a^D156R^-VPR presents observable fluorescence at 25 °C unlike the original system (Figure S15). In all microscopy images mycelia were observed under brightfield (BF) and mCherry filter. Scale bar 100 μm. **b.** *A. nidulans* growth temperature of 25°C is limiting for CRISPR/dLbCas12a-VPR mediated *micA* activation (purple), as no increments in microperfuranone (**1**) are observed. The variant dLbCas12a^D156R^-VPR (green) demonstrated CRISPRa activity at 25°C, achieving a ~4.5-fold increase in the production of **1** compared to the no crRNA control. However, the activity of dLbCas12a^D156R^-VPR was limited compared to the original dLbCas12a-VPR system at 37 °C. **c.** Activity at the alternative PAM site TTCN is increased by the dLbCas12a^D156R^-VPR variant. In all figures, calculated titer (mg L^−1^) values are the mean of three biological replicates whose specific values are indicated as dots, error bars represent SD. Two-sided Welch’s T-test with was performed. Asterisk indicates corrected P-value<0.05. Individual P-values are listed in Table S9.

The presence of protospacer adjacent motif (PAM) near the target is a critical requirement for CRISPR systems, in the case of LbCas12a systems the sequence TTTV. ^52^ The variant LbCas12a^D156R^ has been reported to exhibit improved recognition of the non-canonical PAM sequence TTCN. ^53^ We tested a poly-crRNA array targeting TTCN PAM sites in the 5′ UTR of *micA*, and observed an improved activation mediated by the dLbCas12a^D156R^ variant over the original dLbCas12a system at 37 °C (Figure 4c). In most *A. nidulans* BGC genes around ten canonical TTTV PAM sites can be found in a targetable window for activation (Figure S17). Nevertheless, targeting TTCN can be considered if PAM site availability is a limiting factor.

## Conclusion

In this work, we reported the first application of CRISPRa in an ascomycetous filamentous fungus belonging to the Pezizomycotina taxon, known to harbor diverse BGCs. We demonstrated that CRISPR/dLbCas12a-VPR-mediated multigene activation of BGCs in *A. nidulans* can be a useful tool for discovery of bioactive secondary metabolites. This approach has the potential to be highly scalable due to ease of crRNA array assembly, paving the way to higher throughput natural product genome mining approaches. The CRISPR/dLbCas12a-VPR-based CRISPRa toolkit is built with well-established fungal genetic parts. Hence, along with the generation of a dLbCas12a^D156R^ variant for cultures at room temperature, the CRISPRa toolkit developed here should be transferrable to other filamentous fungi.

Targeting more genes in future works will help understand the broader applicability of CRISPRa and advance guidelines for achieving robust gene activation in filamentous fungi. CRISPRa systems are rapidly evolving and these innovations could be used to upgrade the current fungal CRISPRa system. In particular, some next generation activators have been demonstrated to outperform the activation strength of dCas-VPR systems.^54,55^ Putative activation domains of fungal BGC-specific TFs could also be tested to expand the CRISPRa toolkit. With better guidelines for Cas12a-associated crRNAs design becoming available,^56^ it is possible to further improve fungal CRISPRa efficiency by optimizing crRNA selection and design.

In conclusion, this work represents a valuable expansion to the fungal CRISPR toolbox and provides a foundation for the further development of CRISPRa as a tool for the discovery of novel bioactive fungal natural products via targeted BGC activation. The fungal CRISPRa toolkit also has the potential to be used for elucidation and engineering of natural product biosynthesis by targeting different combinations of biosynthetic genes, for simultaneous activation of multiple pathway-specific TFs, as well as other synthetic biology applications.

## Materials and Methods

### Vector Construction

Main vectors and maps for both Cas12a and Cas9-driven systems are deposited in Addgene with the Addgene Codes indicated in Table S4. All vectors along with their description and the cloning method used in their construction can be found in Table S4. Vectors were generated using one of the following methods: restriction enzyme cloning with PacI, NotI and T4 DNA ligase (New England Biolabs, MA, USA); Type IIS assembly with BsmbI (New England Biolabs, MA, USA) and annealed oligo cloning; *in vivo* homologous recombination in *Saccharomyces cerevisiae* BJ5464 or isothermal assembly with NEBuilder HiFi DNA Assembly Master Mix (New England Biolabs, MA, USA). In all cloning procedures we used *E. coli* 10-Beta electrocompetent (New England Biolabs, MA, USA) or in-lab made equivalent for increased transformation efficiency. All PCR primers and other oligonucleotides used are listed in Table S8, along with their destination construct and source of DNA template if applicable. The AMA1 fungal vector pKW20088^57^, was a gift from Prof Kenji Watanabe, University of Shizuoka, and the vectors pYFAC-riboB, pYFAC-pyroA, pYFAC-CH2, pYFAC-CH3, pYFAC-CH4 were built previously^16^. In all cases when amplifying *P_gpdA_* and *T_trpC_* consisted of the versions present in the expression cassette from pBARGPE1^35^, (obtained from the Fungal Genetics Stock Centre) or the modified version pBARGPE1-LIC^58^. To clone dSpCas9-VPR it was amplified from pAG414GPD-dCas9-VPR^19^, which was a gift from George Church (Addgene plasmid #63801). The coding sequence of dLbCas12a (mutation D832A) was amplified from a plasmid kindly provided by Christian Pflüger which was constructed from pY027^59^, a gift from Feng Zhang (Addgene plasmid # 84742), by site-directed mutagenesis and fused to VPR amplified from pAG414GPD-dCas9-VPR^19^. The sequence of *P_elcA_* was amplified from pYFAC-CH6^16^. The coding sequence of *mCherry* was amplified from pMP7601 ^60^ which was a gift from Alex Andrianopoulos (University of Melbourne). *A. nidulans* sequences were PCR amplified from *A. nidulans* LO8030 gDNA (chromosomal coordinates indicated in Table S5)^61^. *AfP_U3_* was amplified from *Aspergillus fumigatus* 293 gDNA. An adapted version of pGEM-T (Promega) was used to build the Step 1 crRNA and sgRNA cloning vector. The Cas9 sgRNA cloning cassette was synthesized as gBlock and re amplified when fused to *AfP_U3_* (Sequence at Note S1). The *P_gpdA_* Cas12a crRNA cloning cassette was created by annealed oligo cloning (Sequence at Note S1). BsmbI domesticated one-step-cloning vector pCRI008 was built by PCR site directed mutagenesis of PYFAC-pyroA parts (Figure S4a).

### Design and cloning of sgRNA and crRNA

The target sequences of each crRNA or sgRNA are listed in Table S6, along with the corresponding PAM sequence and the distances to target gene start codon and TSS when available^61–63^. The spacers were also verified to pass the bioinformatic off-target test against *A. nidulans* FGSCA4 genome sequence with EuPaGDT^64^. All crRNA and sgRNA were synthesized as oligonucleotides with overhangs whose sequence is indicated in Table S7. Oligos were mixed in equal proportion (10nM), annealed on a thermocycler, if necessary (crRNA arrays) also phosphorylated with T4 polynucleotide kinase (New England Biolabs, MA, USA) and ligated with T4 DNA ligase in previously BsmbI digested vectors.

For some Cas12a crRNA, one-step cloning was possible in the fungal crRNA expression vector pCRI008. The rest of the crRNA were cloned by a 2-vector cloning procedure (Figure S4c). In that case, oligos with encoded crRNA were first cloned into the pGEM-T derived vector pCRI007, and the expression cassette further PCR amplified with primers that reconstituted full *P_gpdA_* and added homology arms. The amplicon was then cloned to the final YFAC fungal vector with homology-based cloning.

For Cas9 sgRNA, the sgRNAs were first cloned into pCRI010, and the PacI NotI flanked expression cassette digested, gel purified and ligated to a PacI NotI digested pCRI011 reporter vector.

### *A. nidulans* strains construction and transformation

*A. nidulans* strains with either dSpCas9-VPR or dLbCas12a-VPR chromosomal expression cassettes were created by polyethylene glycol (PEG)-calcium-based transformation as in Lim et al.^65^ with the previously NotI linearized vectors pCRI001–3 (Table S5) containing 1kb homology regions to facilitate homologous recombination in *A. nidulans 8030* stcJΔ locus. The fragment also contained the *Bar* marker, and colonies were selected for resistance to glufosinate extracted from Basta (Bayer, Vic., Australia) as in Chooi et al.^33^ an the event was confirmed by diagnostic PCR. Complete genotype of the parental strains is indicated in Table S4.

For each transformant strain genotype of the protoplasts used and vectors transformed are listed in Table S4, indicating whether the auxotrophies are complemented by the vectors or supplemented in the media. Protoplasts of *A. nidulans* LO8030 and dCas-VPR expressing parental strains were prepared from germlings as in Lim et al.^65^, mixed with a quarter volume PEG 60% to a final concentration of 10^8^ protoplasts per mL and frozen at −80 °C for later use. AMA1-vectors were transformed into *A. nidulans* protoplasts modifying Lim et al. ^65^ in order to minimize the required transformation volume. In a 2 mL microcentrifuge tube, 60 μL of thawed protoplast solution was incubated with 50 μL of STC buffer (1.2 M sorbitol, 10 mM CaCl_2_, 10 mM Tris–HCl, pH 7.5) and 3 μg of each vector contained in maximum total volume of 10 μL. After 20 min of incubation on ice, 350 μL of the calcium PEG 60% mix was added and mixed gently by inversion, followed by a 20 min incubation at room temperature. After adding 1 mL of STC buffer the mix was spread on Sorbitol Minimal Medium (SMM), that were then incubated for three days at 37 °C to generate transformant colonies.

### Fluorescence microscopy

For each transformant strain, spores from three individual colonies were picked as biological replicates and grown in small petri dishes containing liquid Glucose Minimal Media, GMM, ^16^ to obtain mycelia. Samples were incubated at 37° C and grown overnight, unless other incubation temperature specified. For the samples incubated at 30 °C mycelia were collected after overnight growth, while samples incubated at 25 °C were grown for two days in order to harvest comparable mycelial growth. Fluorescence images were captured on the epifluorescence inverted microscope Eclipse Ti2 (Nikon), using Plan Apo λ 10x /0.45 numerical aperture (NA) objective lens (Nikon) and a Camera DS-Qi2 (Nikon) controlled by NIS Elements Advanced Research (Nikon). Fluorescent microscopy was carried out under a mCherry filter set (562/40 nm excitation, 593 nm dichroic beamsplitter, and 641/75 nm emission), using an 800 ms exposure and 9.6x analog gain unless specified otherwise. Brightfield images were captured at a 300 ms exposure time with 1x analog gain. Images were recorded using NIS-Elements Advanced Research software package (Nikon).

### Culture conditions and crude extract preparation

For each transformant strain, spores from three individual colonies were picked as biological replicates for culture analysis and re-streaked individually in a solidified GMM plate to be cultivated for three days at 37 °C. Spores were harvested from plates in 1 mL of 0.1% Tween 80 (Sigma, MO, USA) and after counting under Neubauer chamber, 2×10^8^ spores were inoculated into 250-mL flasks containing 50 mL liquid GMM medium as described previously. ^16^ Additionally, ampicillin was added to 50 μg mL^−1^ and riboflavin, pyridoxine, uracil and uridine was supplemented, if necessary, as indicated in Table S4. Cultures were grown for 2.5 days with shaking set to 200 rpm and 37 °C, unless other temperature indicated. In the case only of the samples needing *P_alcA_* promoter induction, cyclopentanone at a final concentration of 10 mM was added to the medium after 18 h of incubation. At the end of the culture, 20 mL of media was collected in 50-mL Falcon tubes by filtration with Miracloth (Milipore, MA, USA). The metabolites were extracted from the liquid culture with 20 mL of an organic solvent mixture containing ethyl acetate, methanol and acetic acid (89.5:10:0.5 ratio). The crude extracts were dried down *in vacuo* and re-dissolved in 0.3 mL of methanol for LC-DAD-MS analysis.

### Metabolic profile analysis by LC-DAD-MS

The analyses of the metabolite profiles were performed on an Agilent 1260 liquid chromatography (LC) system coupled to a diode array detector (DAD) and an Agilent 6130 Quadrupole mass spectrometer (MS) with an electrospray ionization (ESI) source. In all cases 3 μL of the methanol dissolved crude extract was injected. Chromatographic separation was performed at 40 °C using a Kinetex C18 column (2.6 μm, 2.1 mm i.d. 3 100 mm; Phenomenex). Chromatographic separation was achieved with a linear gradient of 5–95% acetonitrile-water (containing 0.1% v/v formic acid) in 10 minutes followed by 95% acetonitrile for 3 minutes, with a flow rate of 0.70 mL min^−1^. For the multiple target CRISPRa experiments, the gradient was extended to 20 min for better separation. The MS data were collected in the m/z range 100– 1000 in positive ion mode and UV observed at DAD λ=254.0±4.0 nm.

Peak areas were determined by peak integration of DAD λ=254 nm chromatogram using Masshunter Workstation Qualitative Analysis (Agilent). To quantify microperfuranone (**1**) samples were compared to a calibration curve. To this end, a standard of **1** was prepared by weighing approximately 14 mg of purified **1** and diluting in methanol. Several dilutions of the standard were measured by LC-DAD-MS. The standard was weighted, and the procedure repeated independently three times (Figure S20). A representative regression fit to zero was used to quantify **1**, consisting in different concentrations of **1** (0.6, 0.5, 0.4, 0.3, 0.2, 0.1, 0.05, 0.01, 0.001, 0 mg L^−1^) with three injection replicates (Figure S3b). The linear regression coefficient was used to extrapolate the concentrations in the crude extract to the culture media concentrations.

For the calculated additive of single crRNA mediated production of **1** in Figure 2e, the negative control mean was added to the summation of the difference between the mean of each individual crRNA production and the negative control mean, for the four crRNAs tested.

### LC-MS/MS analysis

Selected samples were analyzed by LC-MS/MS on a Thermo Scientific Fusion Orbitrap coupled to a Thermo Ultimate 3000 UHPLC. The column used was an Agilent Poroshell 120 SB-C18 (2.1 × 30 mm, 2.7 μm) with a 20 min linear gradient of 5–95% acetonitrile-water containing 0.1% v/v formic acid. Precursor ion data was collected for m/z 200 to 300 in positive ion mode. Fragmentation was achieved with the higher-energy collisional dissociation cell set to a collision energy of 15. Fragment identification was aided by CFM-ID^43^ predictions based on hypothesized structures.

### Compound isolation and NMR structural characterization

For the microperfuranone (**1**) standard purification, 2 L of 2-days culture media (post-induction) from strain 52 (Table S4), was extracted with a mix of ethyl acetate, methanol and acetic acid (89.5:10:0.5). The crude extract was dried *in vacuo*, resuspended in methanol and loaded onto a Sephadex LH-20 (GE Healthcare, Little Chalfont, UK) column for fractionation. Fractions containing the target compound were combined and further purified by semi-prep HPLC with a C18 column (Grace, 5 μm, 10 × 250 mm) (isocratic, 40% acetonitrile-water, 4.3mL min^−1^).

For purification of dehydromicroperfuranone (**2**), 4 L of 2-days culture media (post-induction) from strain 54 (Table S4), after induction, was loaded onto a customized Diaion HP-20 (Sigma, PA, USA) column pre-equilibrated with water. The column was then flushed with 2 column-volume of water and eluted with methanol. The eluent was dried *in vacuo*. The crude extract was resuspended with methanol and fractionated using the Sephadex LH-20 column. Fractions containing the target peaks were combined and further purified by a Reveleris flash chromatography system (Grace) using a C18 Preparative column (Agilent, 5 μm, 21.2 × 150 mm). A gradient method (55% acetonitrile-water to 85% acetonitrile-water in 12 mins, 10 mL min^−1^) was applied for the separation of peak III with peaks I-III.

For structural characterization of **1** and **2**, nuclear magnetic resonance (NMR) spectra were collected on Bruker Avance IIIHD 500/600MHz NMR spectrometers, with either CDCl_3_-*d* or MeOD-*d_4_* as solvents. NMR data in CDCl_3_ was in good agreement with the published data (Table S2; Table S3, Figures S18-19).

### Computational analysis of *A. nidulans* features

All Python code used are implemented in a Jupyter notebook which is available at https://github.com/gamcil/5_UTR_analysis/ alongside accompanying data. The genome assembly and corresponding gene annotations for *A. nidulans* FGSC A4 were obtained from FungiDB (denoted version 46)^63,66^. Coordinates of predicted BGCs were obtained from the *A. nidulans* portal on the Joint Genome Institute’s MycoCosm resource^67^. The genome was parsed for gene features using Python scripts. Genes falling within cluster boundaries were grouped, forming an additional dataset to facilitate comparison between the whole genome and BGC genes.

Lengths of 5′ UTRs were determined for all genes and BGC genes, based on features when available, filtering the genes with 5′ UTRs equal zero. Histograms for each dataset were plotted using the Matplotlib library.

To determine the frequency of Cas9 and Cas12a PAM sites, upstream regions for all genes were isolated by taking up to 400 bp upstream of the start codon. When intergenic distance was less than 400 bp, the distance between the start codon and the end of the previous gene was used. Frequencies of the different PAM sites were obtained through regular expression searches of the PAM sequences considering both strands. Histograms for each dataset were plotted as mentioned above.

### Alignment of Cas12a structures for residue identification

Crystal structures of LbCas12a (PDB ID: 5XUS) ^50^ and AsCas12a (PDB ID: 5B43) ^49^ were aligned with PyMOL (The PyMOL Molecular Graphics System, Version 1.7.6.3) in order to facilitate identification of the equivalent residue to AsCas12a E174.

### Statistical analysis

Statistical analysis was done using GraphPad Prism 8.3.0. All data were analyzed with three biological replicates and Two-sided Welch’s T-test with Holm-Šídák multiplicity correction per figure, using an alpha of 0.05. All Welch’s T-test P-values calculated for each experiment along the details for the multiplicity adjustment are found in Table S9. Fisher’s exact test two-sided was also performed with GraphPad.

## Supporting Information

**Supplementary File 1** (PDF) Figures S1–20. Fluorescence microscopy images with biological replicates, bioinformatic analysis of *Aspergillus nidulans* 5′ UTR features and PAM site frequency, quantification of additional *micA* CRISPRa assays, complete chromatograms of multiple gene activation of *mic* cluster with quantification, dehydromicroperfuranone isolation chromatograms, dehydromicroperfuranone NMR spectra, Cas12a protein alignment, microperfuranone standard curve and NMR spectra, overview of one-step cloning of Cas12a crRNA. Note S1 with sequence of crRNA/sgRNA expression cassettes cloning sites. Tables S1–3 containing NMR and LC/MS-MS analysis summary.

**Supplementary File 2** (Excel) Additional Tables S4–9 are found as worksheet tabs in a single Excel file. It contains: strains used in this study, vectors used in this study, protospacers targeted, oligonucleotides used to create crRNA/sgRNA, oligonucleotides, statistical analysis.

## Data Availability

All data generated or analyzed during this study are included in the manuscript and supporting files. All Python code used in these analyses is implemented in a Jupyter notebook which is available at https://github.com/gamcil/5_UTR_analysis/ alongside accompanying data. Indicated plasmid maps are available in Addgene.

## Author contribution

I.R and Y.H.C conceived the project. I.R., C.W. and Y.H.C. wrote the manuscript. I.R. designed the constructs. I.R., C.W. and R.W. contributed to cloning. I.R. and C.W. performed the experiments and the data analysis. J.H. performed the NMR structural elucidation. C.L.M.G. wrote the code for computational analysis.

## Acknowledgements

Y.H.C. and this project is supported by an ARC Future Fellowship (FT160100233). I.R. is recipient of an UWA PhD Scholarship, J.H. and C.L.M.G. on Australian Government Research Training Program Scholarships. NMR and LC-MS/MS were performed at the UWA Centre for Microscopy, Characterisation and Analysis (CMCA). We thank Berl Oakley for *A. nidulans* LO8030 strain, and Christian Pflüger and Andrew Piggott for helpful discussion.

## Abbreviations

5′ UTR: 5′ Untranslated Region
AMA1: Autonomous maintenance in *Aspergillus*
BGC: Biosynthetic Gene Cluster
CRISPR: Clustered Regularly Interspaced Palindromic Repeats
CRISPRa: CRISPR activation
crRNA: CRISPR RNA
dCas: DNAse deactivated CRISPR associated protein
dLbCas12a: dCas from *Lachnospiraceae bacterium*
dSpCas9: dCas from *Streptococcus pyogenes*
MD: micA Downstream TSS
MU: micA Upstream TSS
NRPS-like: nonribosomal peptide synthetase-like
PAM: Protospacer Adjacent Motif
PDB: RCSB Protein Data Bank ID
RNAP: RNA polymerase
sgRNA: single guide RNA
VPR: Tripartite activator VP64-p65-Rta
TSS: Transcription Start Site
TF: transcription factor
SM: secondary metabolite

