## Supplementary Information File 1 for "CRISPR-mediated activation of biosynthetic gene clusters for bioactive molecule discovery in filamentous fungi"

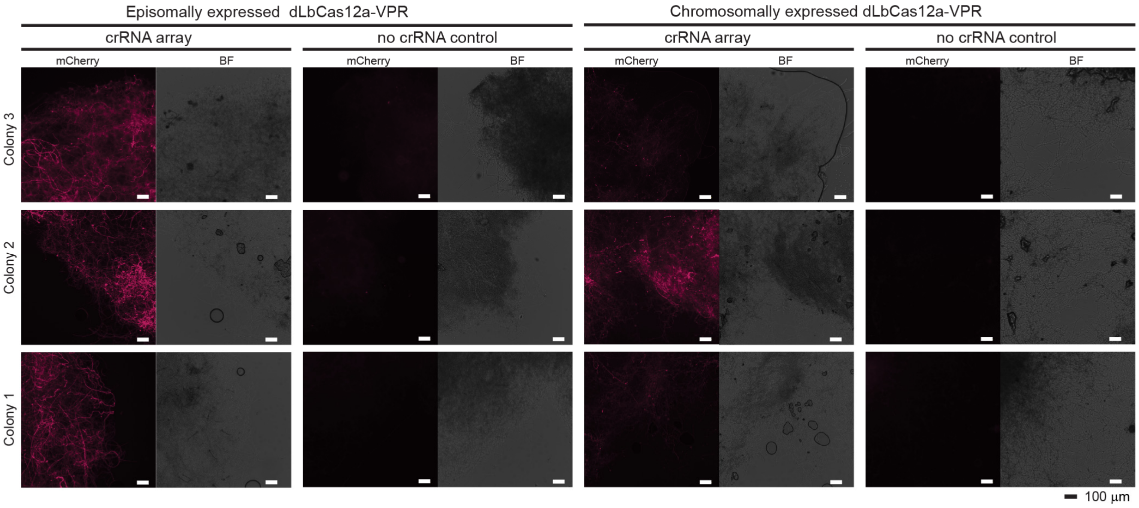


**Figure S1:** CRISPR/dLbCas12a-VPR proof-of-concept replicates. Fluorescence microscopy images of *A. nidulans* mycelia demonstrate mCherry fluorescence when the poly-crRNA array is present in both episomally and chromosomally expressed dLbCas12a-VPR systems, distinct from the no crRNA control. This is indicative of activation of the fluorescent reporter $P_{elcA}$-*mCherry* by CRISPR/dLbCas12a-VPR. The spores for each sample were collected from three individual colonies and grown overnight in liquid stationary culture at 37 °C. Samples with similar mycelial growth were observed under mCherry filter and brightfield (BF). Scale bar 100 $\mu$m.

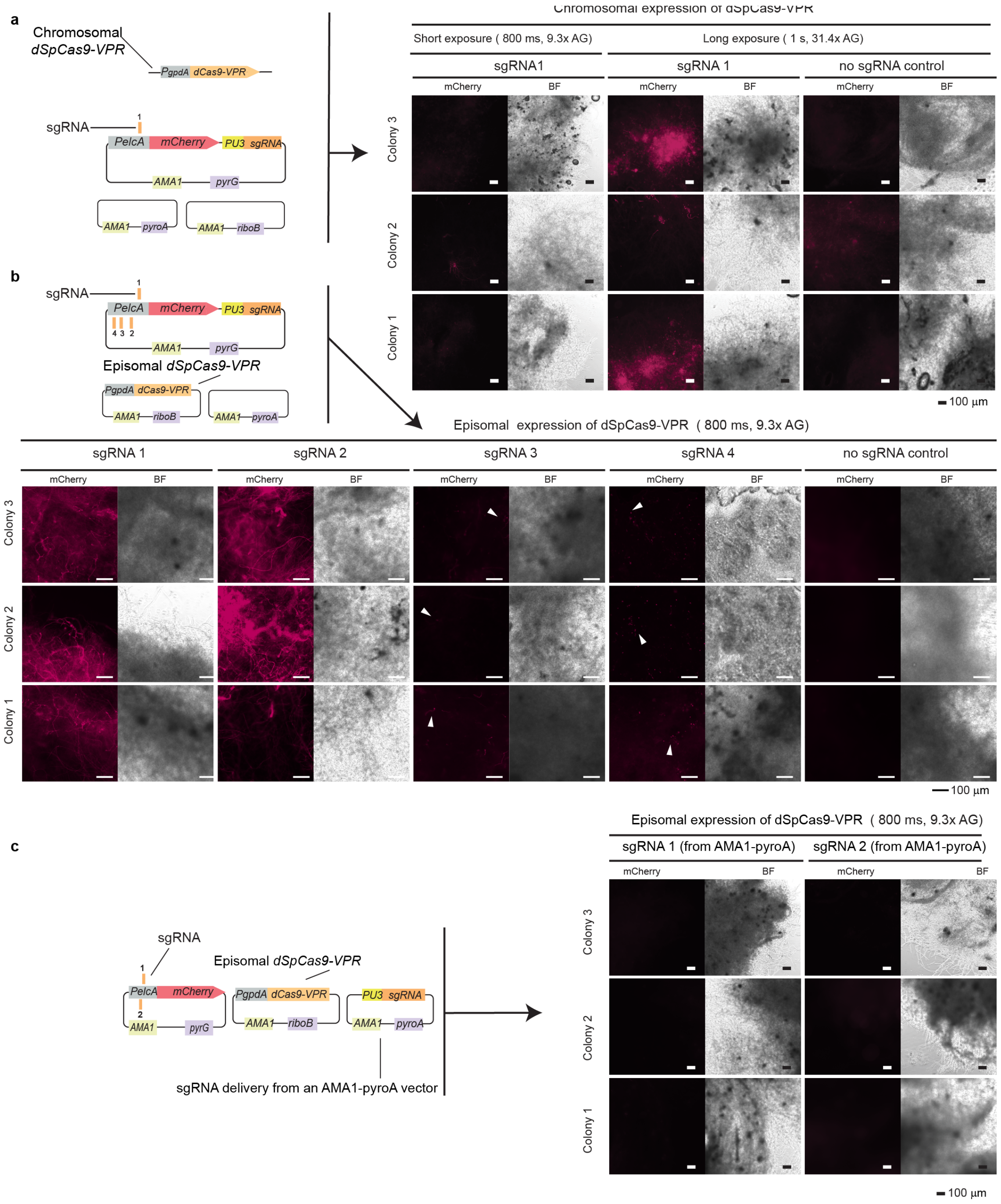


**Figure S2.** CRISPR/dSpCas9-VPR mediated activation of $P_{elcA}$-*mCherry* is dependent on CRISPRa component delivery and sgRNA targeting position.**a.** CRISPRa samples with dSpCas9-VPR chromosomally expressed and sgRNA 1 (represented in diagram) resulted in low activation the mCherry reporter, although samples were distinct to the no sgRNA control when observed at prolonged exposure times and increased sensitivity (1s exposure and 31.4x analog gain). **b.** AMA1-encoded CRISPR/dSpCas9-VPR system (represented in diagram)

resulted in strong fluorescence observable in all mycelia at short exposure times when targeting the reporter construct with sgRNA 1 or sgRNA 2. The two sgRNA targeting regions further away from the gene start codon (sgRNA 3 and sgRNA 4) resulted in low fluorescence localized in spores or isolated mycelia, but distinguishable from the no sgRNA negative control. Images are 2× digitally zoomed to show the fluorescence of the spores (white arrows). **c.** The strong activation by the episomally delivered dSpCas9-VPR system observed with sgRNA 1 and sgRNA 2 is abolished when the sgRNA is delivered encoded in a separate AMA1-pyro vector (represented in adjacent diagram). The spores for each sample were collected from three individual colonies and grown overnight in liquid stationary culture at 37 °C. Samples with similar mycelial growth were observed under mCherry filter and brightfield (BF). Scale bar 100 μm.

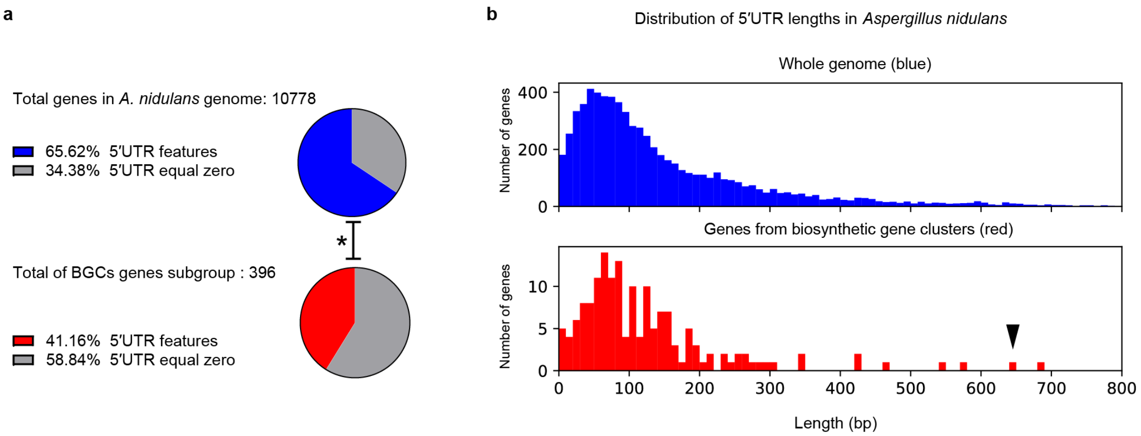


**Figure S3.** *Aspergillus nidulans* 5′ UTR features and annotation availability. **a.** Access to transcription starting site (TSS) information for CRISPRa targeting is limited in the current version of the *A. nidulans* genome. The proportion of genes that have annotated 5′ UTR lengths greater than zero is smaller for BGCs compared to the whole genome. This could be due to the low expression of most BGCs genes in the growth conditions in which the transcriptomic data was acquired [1]. Asterisk represents the significant result of Fisher's exact test two-sided p<0.0001. **b.** Distribution of 5′ UTR length in *Aspergillus nidulans* genes for the whole genome (blue) and in genes that fall within BGC boundaries only (red), considering only genes with 5′ UTR annotation bigger than zero. Arrow indicates the *micA* 5′ UTR length.

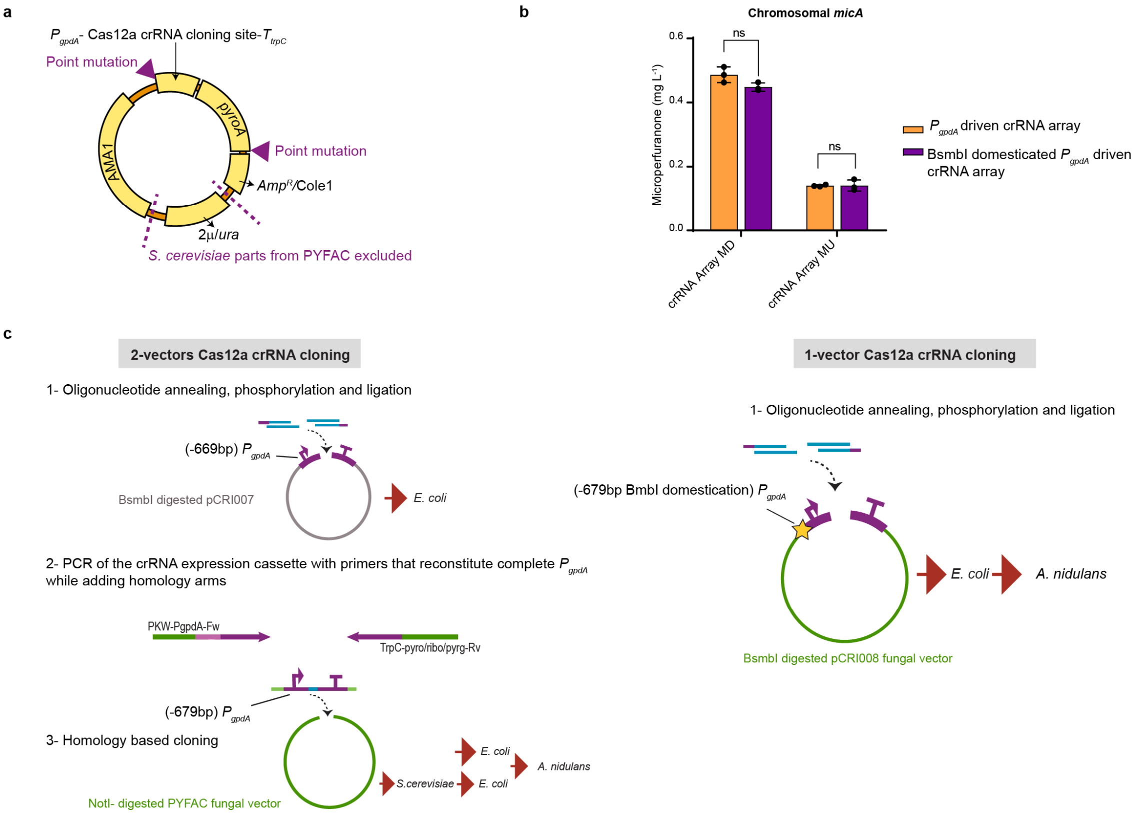


**Figure S4.** One-step cloning of Cas12a crRNA with pCRI008. **a.** Diagram of the BsmbI domestication strategy used to create the vector pCRI008. Two point mutations were introduced, one in $P_{gpdA}$ promoter from the crRNA expression cassette, and the other in the *pyroA* marker terminator region. The components of pYFAC for replication in *Saccharomyces cerevisiae* were eliminated to avoid BsmbI sites. **b.** No difference in microperfuranone (**1**) activation mediated by CRISPRa is observed between systems with crRNA array expression driven by the original $P_{gpdA}$ sequence (orange) and the BsmbI domesticated version of $P_{gpdA}$ (purple), confirming the null effect of the mutation. Titers of **1** (mg $L^{-1}$) values are the mean of three biological replicates, specific values of which are indicated as black dots, bars represent SD. Two-sided Welch's T-test was performed. Asterisk indicates P-value<0.05. Individual P-values are listed in Table S9. **c.** The simplified one-vector cloning strategy with the BsmbI-domesticated pCRI008 versus the original two-vector cloning strategy. In the two-vector strategy, a shorter $P_{gpdA}$ lacking a BsmbI site was cloned into a pGEM-T backbone to create a vector (pCRI007) for Type IIS restriction enzyme cloning. The cloned crRNAs in Pcri007 is later amplified by primers that would reconstitute the full $P_{gpdA}$ sequence when the expression cassette is incorporated into a fungal vector by homology-based cloning.

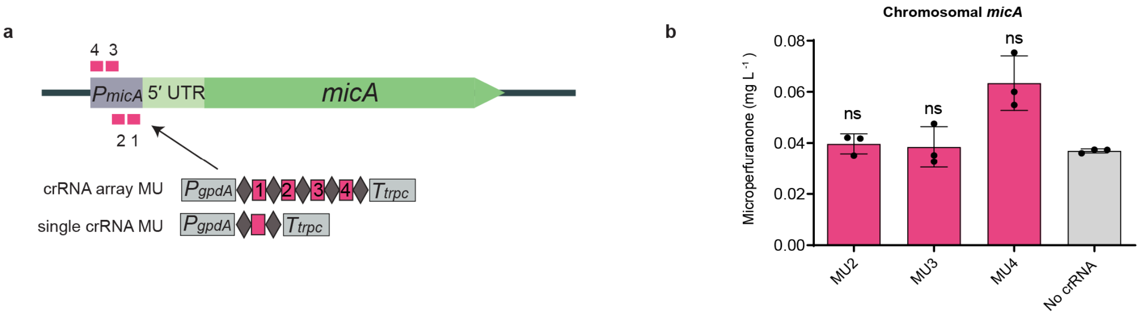


**Figure S5.** Testing of individual MU crRNAs. **a.** Chromosomal *micA* gene scheme with individual crRNA MU (magenta) target sites indicated. Numbers assigned to crRNA MD are indicative of targeting position in respect to *micA* TSS, with their position in the crRNA array MD also indicated in the scheme **b.** No significant changes in the production of microperfuranone are observed targeting with single crRNAs compared to the no crRNA control. Microperfuranone titer (mg $L^{-1}$) values are the mean of three biological replicates in which specific values are indicated as black dots, error bars represent SD. Two-sided Welch's T-test with Holm-Šídák multiplicity correction per figure was performed. Asterisk indicates corrected P-value<0.05. Individual P-values are listed in Table S9.

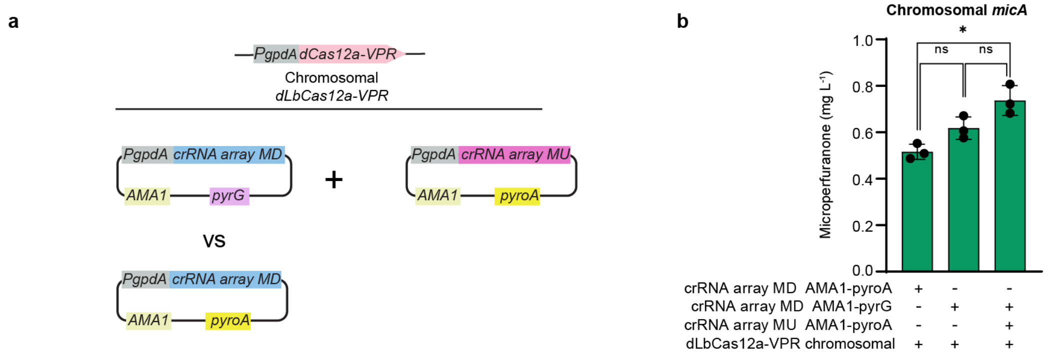


**Figure S6.** CRISPRa of *micA* with both MD and MU crRNA arrays. **a.** Overview of double crRNA array mediated activation of *micA* in strains harboring chromosomally integrated *dLbCas12a-VPR*. In order to allow co-transformation of two crRNA arrays, crRNA array MD was re-cloned into an AMA1-pyrG vector. This allowed comparing the performance of crRNA array MD in both AMA1-pyrG and AMA1-pyroA vectors. **b.** Double crRNA array (8 crRNA) mediated activation of *micA* resulted in microperfuranone titers higher than the four-crRNA array MD alone when delivered from AMA1-pyroA. However, there is no significant increase from the production observed by crRNA array MD alone when delivered from AMA1-pyrG. The differences in the production mediated by crRNA MD when using the markers *pyrG* and *pyroA* is not significant, although it might suggest that the marker used for crRNA delivery could influence the system performance. Microperfuranone titer (mg $L^{-1}$) values are the mean of three biological replicates, of which specific values are indicated as black dots, bars represent SD. Two-sided Welch's T-test with Holm-Šídák multiplicity correction per figure was performed. Asterisk indicates corrected P-value<0.05, (ns) not significant. Individual P-values are listed in Table S9.

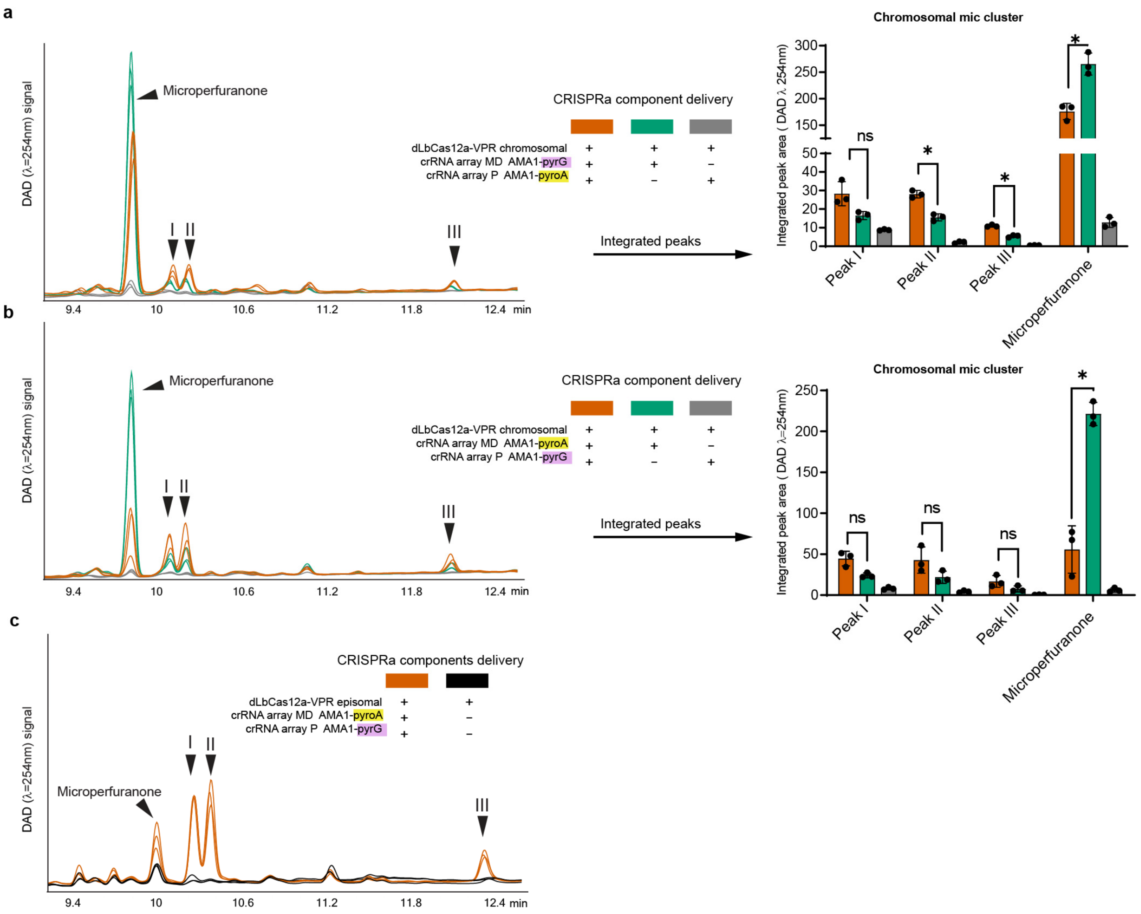


**Figure S7.** Multiple gene activation of *mic* cluster replicates, controls and quantification. **a–b.** Overlaid DAD (λ=254 nm) chromatograms of acidified ethyl acetate mix media crude extracts from strains co-transformed with the multiple crRNA arrays MD and P (orange), crRNA array MD (green) and crRNA array P (grey); marker combination for crRNA array delivery in **a** for multiple targeting (orange), crRNA MD in an AMA1-pyrG vector and crRNA P in a AMA1-pyroA vector; combination in for **b** (orange), crRNA MD in an AMA1-pyroA vector and crRNA P in an AMA1-pyrG vector. Both multiple crRNA arrays (orange) for combinations **a** and **b** showed partial conversion of the precursor microperfuranone to the peaks I–III. Compared to **a**, combination **b** showed a stronger decrease in the titer of the precursor microperfuranone and higher titer of the peaks I–III. **c.** Expression of dLbCas12a-VPR from the multicopy AMA1-vector with the same crRNA array delivery combination as **b** resulted in a further increase of peaks I–III production relative to microperfuranone. Chromatograms in **a**,**b**,**c** are represented at the same scale. In peak area quantification values are the mean of three biological replicates, of which specific values are indicated as black dots, bars represent SD. Two-sided Welch's T-test with Holm-Šídák multiplicity correction per figure was performed. Asterisk indicates corrected P-value<0.05, (ns) not significant. Individual P-values are listed in Table S9.

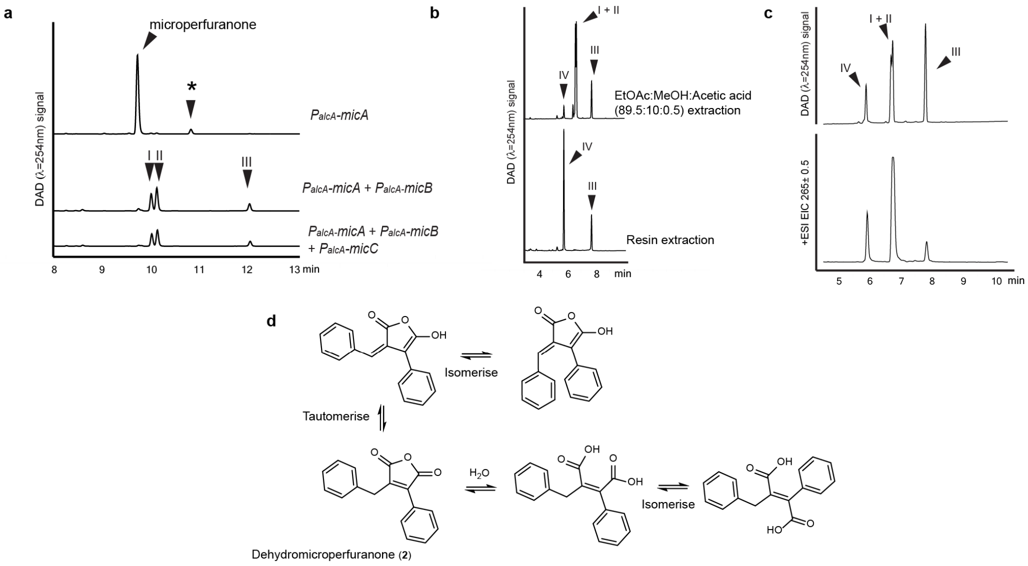


**Figure S8.** Verification of *mic* cluster product by promoter exchange and dehydromicroperfuranone isolation. **a.** DAD (λ=254 nm) chromatograms from *mic* cluster genes (*micA*–*C*) expression from an alcohol inducible promoter ($P_{alcA}$). Overexpression of *micA* results in the expected microperfuranone peak and a smaller unidentified peak likely to be a side product of microperfuranone with m/z 251 (asterisk). Co-expression of *micA* and *micB* resulted in the production of the peaks I–III peaks as expected. Co-expression of *micA*–*C* results in the same metabolic profile of *micA*–*B*, indicating that *micC* is likely not necessary to produce peaks I–III. All chromatograms run with a gradient of 20 min. **b.** Metabolic profile of highly concentrated samples from culture media crude extract obtained when scaling up for purification showing peaks I–III and a smaller IV of identical *m*/*z* 265 when extracted with acidified ethyl acetate and methanol mix. When extracting the same media with a solid resin we observe only the peaks IV and III. All chromatograms with a gradient of 10 min. **c.** LC-

DAD-MS analysis of the purified peak III used for NMR analysis (top, DAD chromatogram λ=254 nm; bottom, extracted ion chromatogram m/z 265). The purified peak III NMR sample in chloroform-*d*, which appeared as a pure single chemical entity based on $^{1}H$ and $^{13}C$ NMR analysis (Table S2, Figures S9–10), was dried and reconstituted in methanol for the LC-DAD-MS analysis. LC-DAD-MS of co-purified peaks I–II shared the same profile. $^{1}H$ NMR analysis of the reconstituted sample in methanol-*d* also showed that the compounds existed as mixed forms (Figures S13–14). **d.** Possible interchangeable forms of dehydromicroperfuranone based on possible tautomerization, interconversion between closed-ring anhydride and open-ring diacid form, and *cis-trans* isomerization of maleic acid to fumaric acid. Note: *m*/z 283 was occasionally observed to coexist with *m*/*z* 265 for the peaks I–IV suggesting that some of the *m*/*z* 265 ions detected could be $[M+H-H_2O]^+$. However, it is difficult to determine which peak belongs to which chemical form, as the isolated peaks appear as a single entity in chloroform-*d* during NMR analysis and converted back to multiple peaks during LC-DAD-MS.

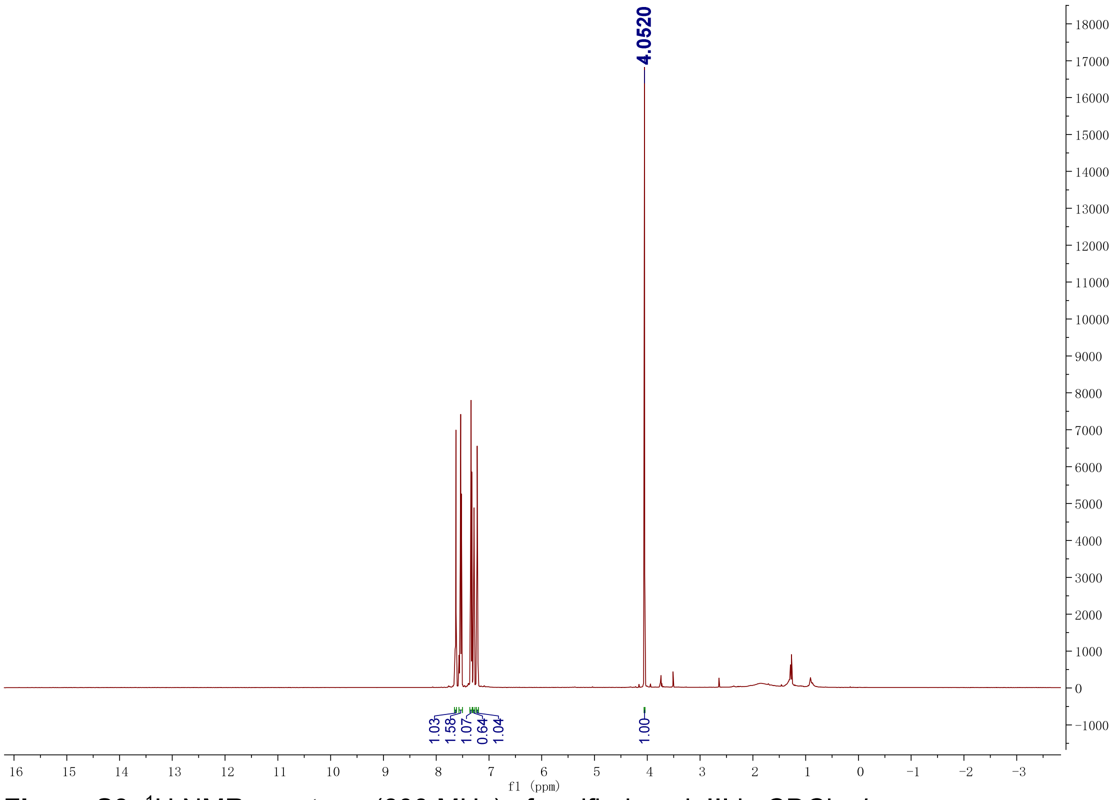


**Figure S9.** $^{1}H$ NMR spectrum (600 MHz) of purified peak III in $CDCl_3$-*d*.

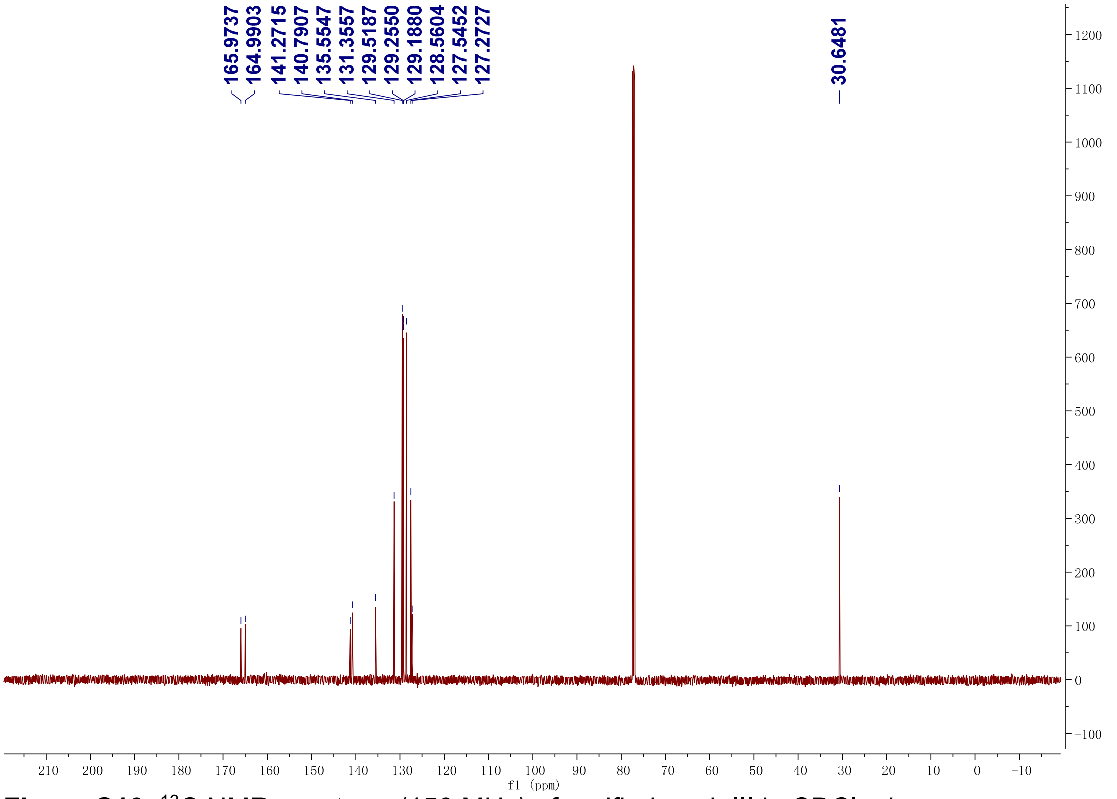


**Figure S10.** $^{13}C$ NMR spectrum (150 MHz) of purified peak III in $CDCl_3$-*d*.

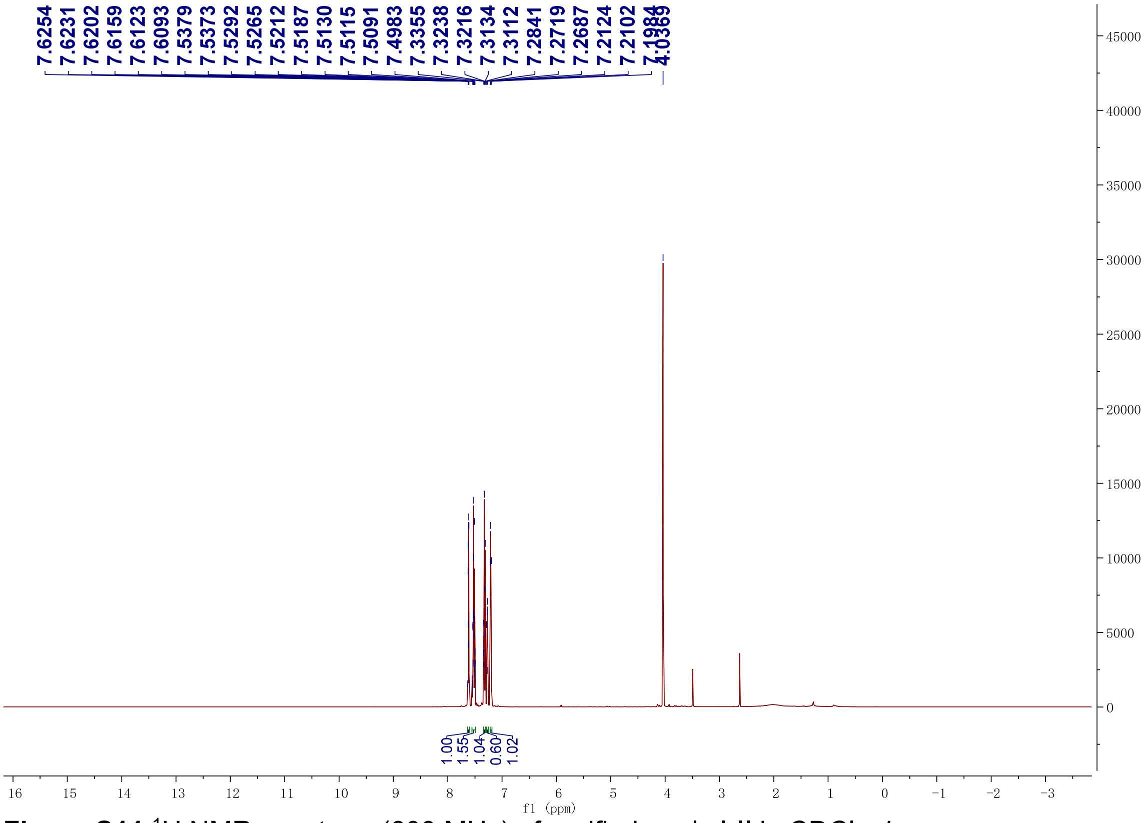


**Figure S11.**[1]H NMR spectrum (600 MHz) of purified peals I-II in $CDCl_3$-*d*.

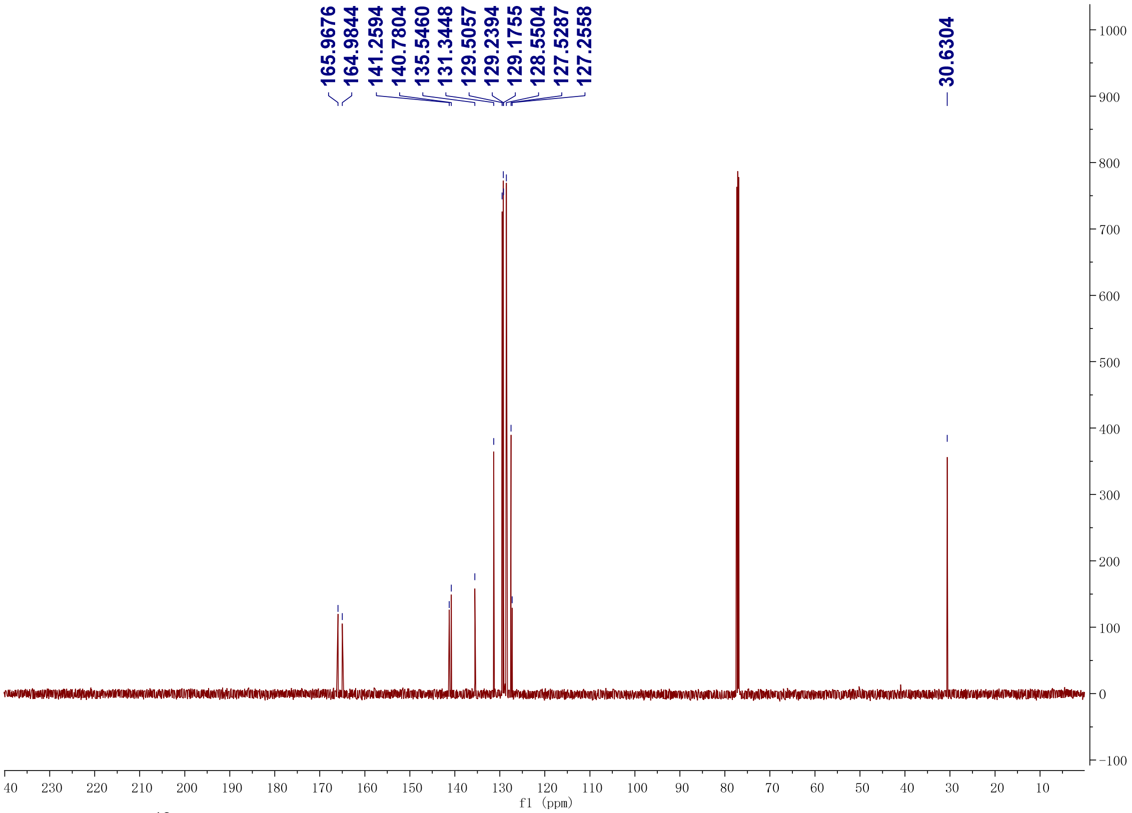


**Figure S12**.[13]C NMR spectrum (150 MHz) of purified peaks I-II in $CDCl_3$-*d*.

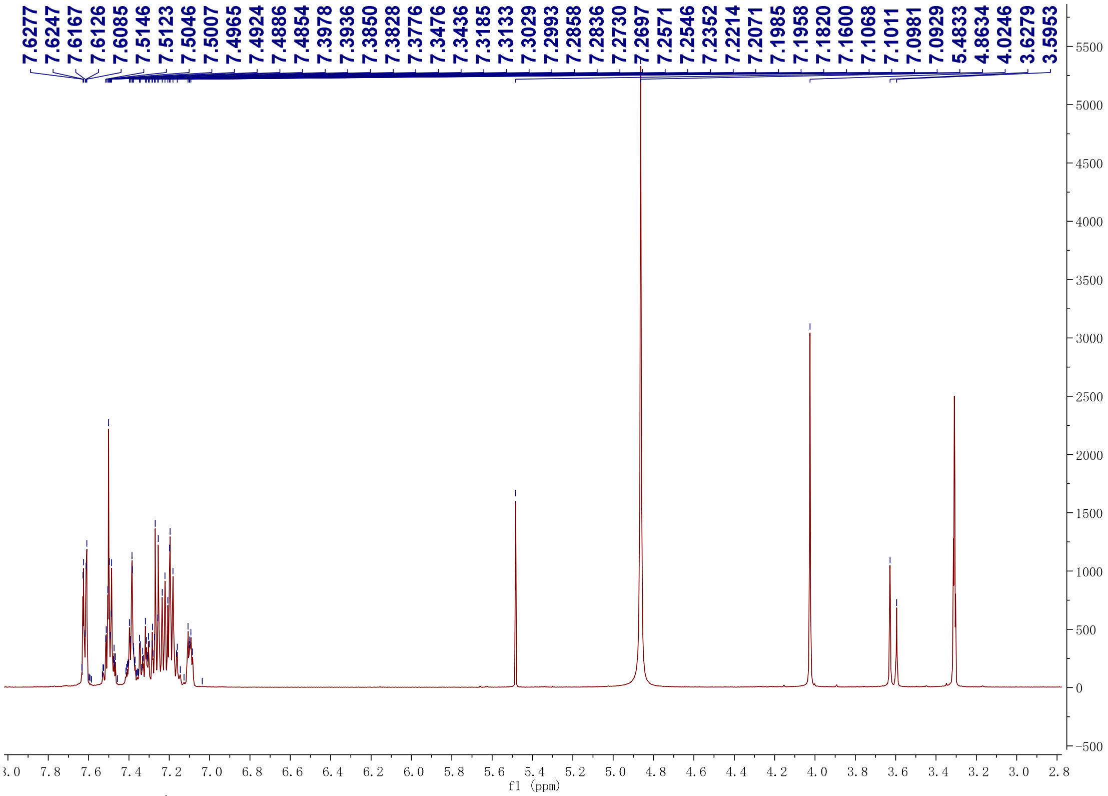


**Figure S13.** $^1$H NMR spectrum (500MHz) of purified peaks **I-II** in MeOD.

.

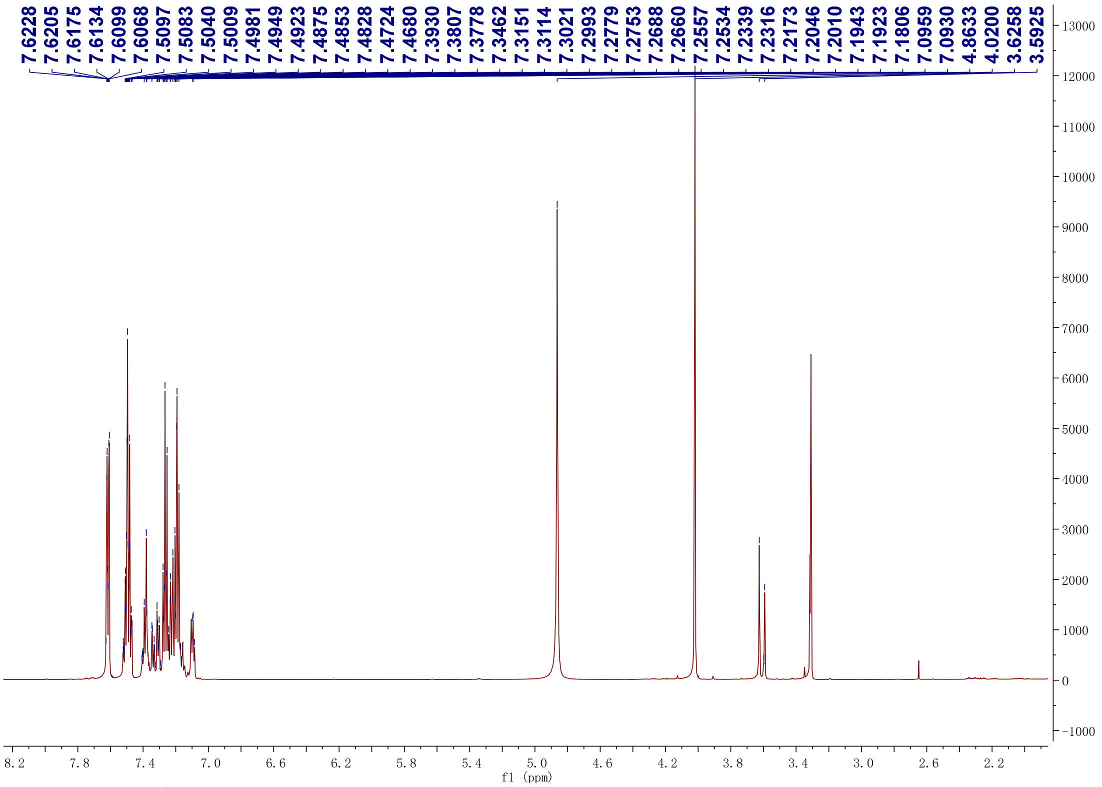


**Figure S14.** $^1$H NMR spectrum (500MHz) of purified peak **III** in MeOD.

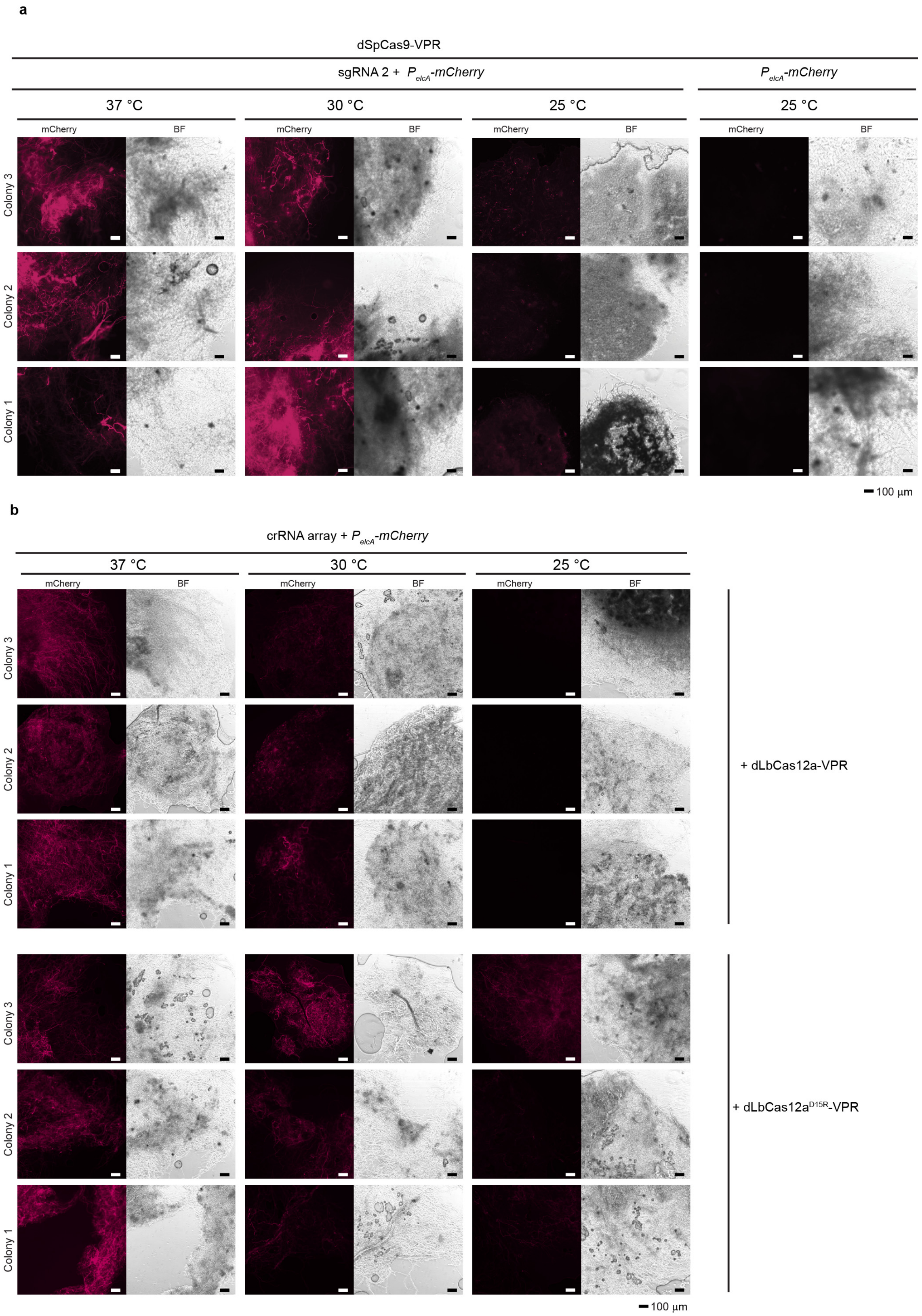


**Figure S15.** CRISPR-mediated activation of $P_{elcA}$-*mCherry* fluorescent reporter is limited at low temperatures. **a**. The activation observed in mycelia by CRISPR/dspCas9-VPR at 37 °C is similar to that observed at 30 °C but lower at 25 °C (room temperature) in mycelia at a stage of equivalent growth to that observed in the other temperature conditions. Fluorescence at 25 °C was still distinguishable from the no sgRNA control. **b**. CRISPR/dLbCas12a-mediated activation was observed for mycelia grown at 37 °C and 30 °C but not observed at 25 °C. The variant dLbCas12a$^{D156R}$-VPR presents observable fluorescence signal at 25 °C unlike the original system. In all cases, the spores for each sample were collected from three individual colonies and grown in liquid stationary culture overnight in the case of 37 °C, and 30 °C and two days for the samples at 25°C. The photos represent the mCherry channel and bright-field (BF). Scale bar 100 $\mu$m.

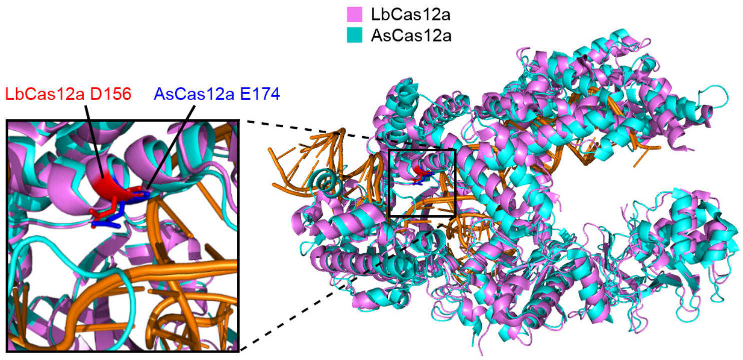


**Figure S16.** LbCas12a and AsCas12a crystal structures alignment for equivalent residue identification. Protein alignment visualized in Pymol of LbCas12a (pink) (PDB ID: 5XUS) [2] and AsCas12a (PDB ID: 5B43) [3] (light blue) crystal structures indicates that residue D156 from LbCas12a (Red) is an equivalent residue to E174 from AsCas12a (blue).

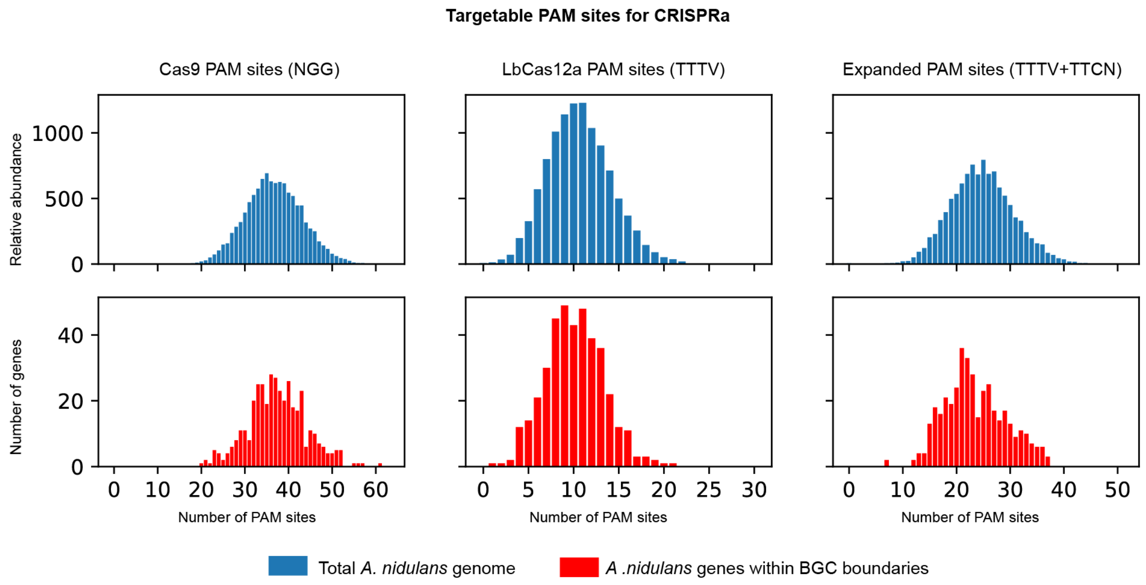


**Figure S17.** Amount of PAM sites identified in the targetable window of *A. nidulans* genes (blue) and genes located in BGCs (red). Targetable window was defined as 400bp upstream of the start of the gene (TSS if available, otherwise -100bp of the start codon) or shorter if intergenic distance is less than 400bp. Even though LbCas12a has fewer targeting sites compared to Cas9, the median is 10 PAM sites per gene. In cases with limiting number of PAM sites, the use of the non-canonical site TTCN with the dLbCas12a$^{D156R}$-VPR variant could be explored, at the cost of losing efficiency for the canonical site, based on our observations at 37 °C (see Figure 4b).

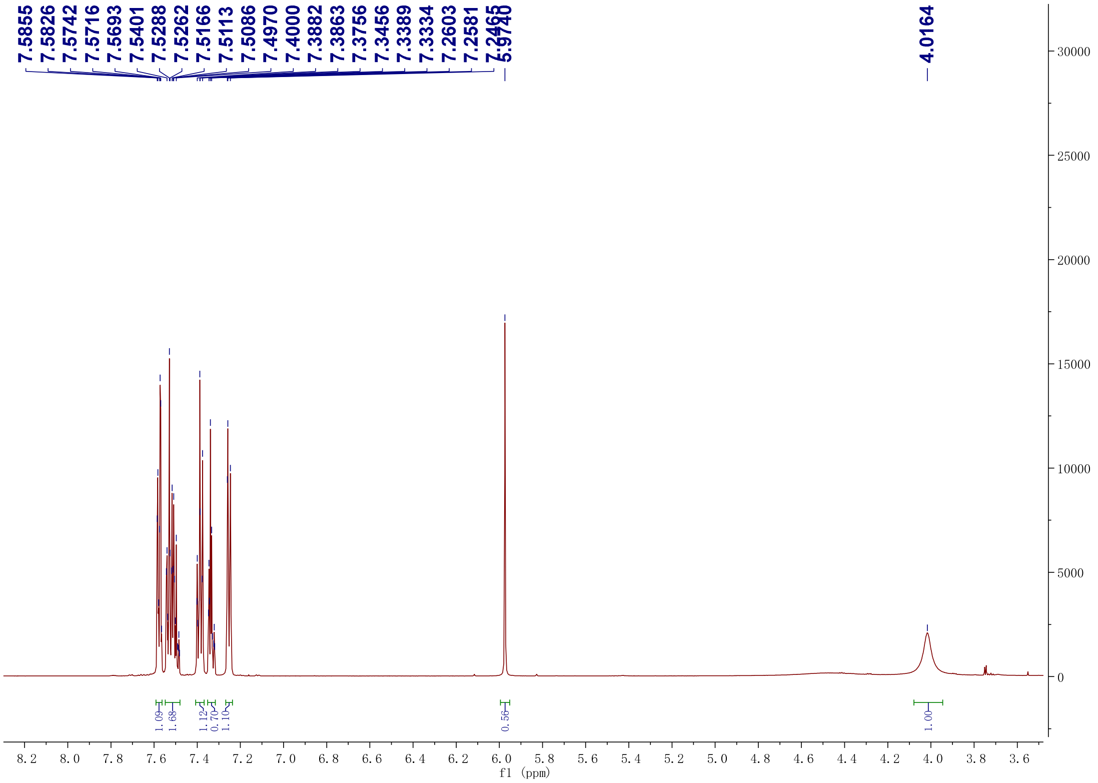


**Figure S18.** [1]H NMR spectrum (600 MHz) of microperfuranone in $CDCl_3$-*d* for standard preparation.

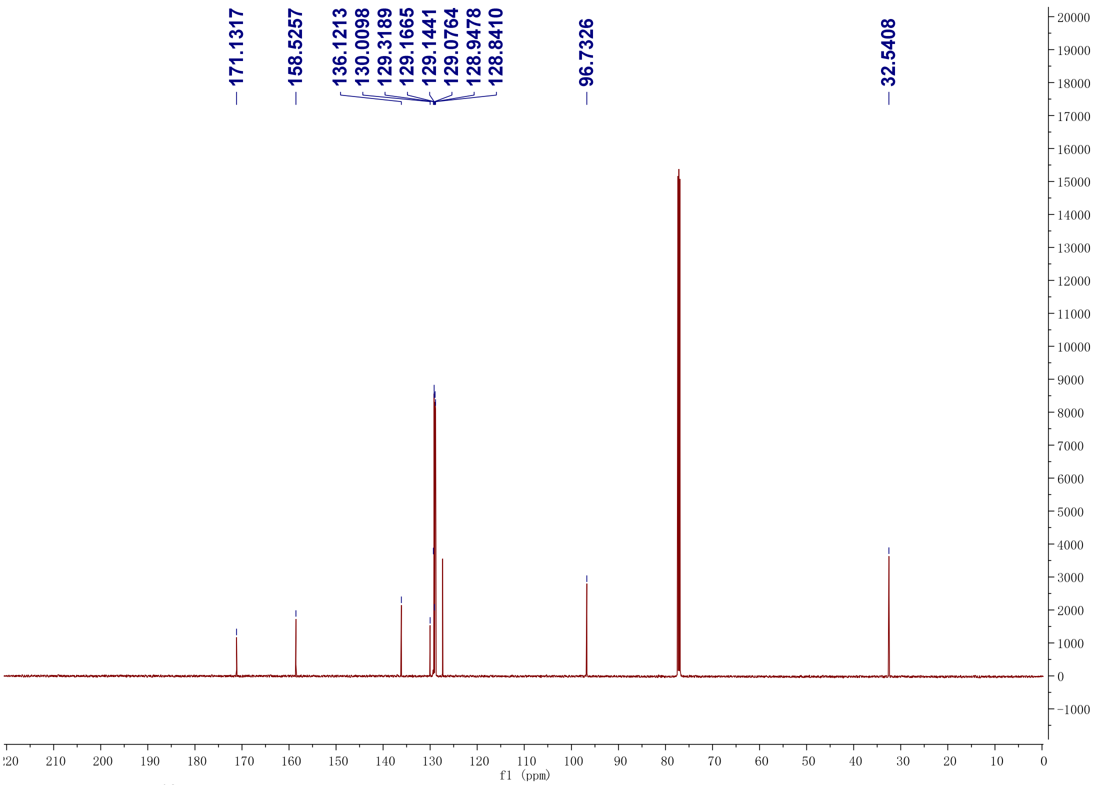


**Figure S19.**$^{13}$C NMR spectrum (150 MHz) of microperfuranone in $CDCl_3$-*d* for standard preparation.

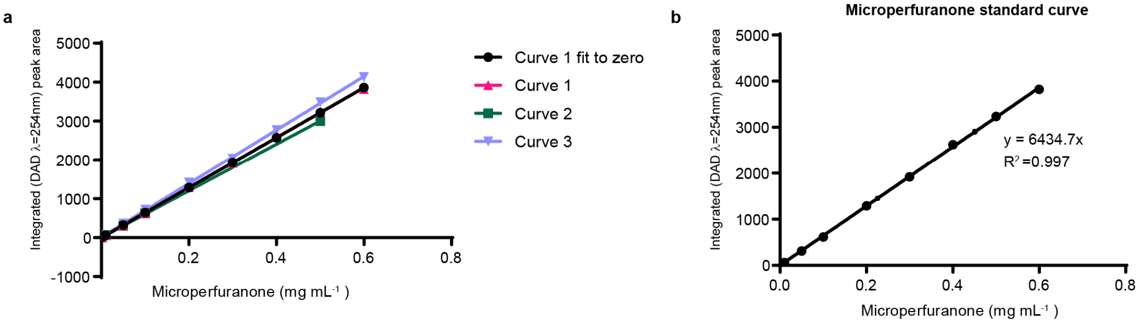


**Figure S20.** Calibration curve for microperfuranone quantification. **a.** Independent microperfuranone standard curves observing peak area by DAD (λ=254 nm) at different concentrations. Individual values indicated as points. **b.** Curve 1 fit to zero was chosen as representative for quantification, points are the mean of three LC-DAD-MS injection technical replicates per point, SD is represented. The equation represents the linear regression used for microperfuranone quantification, where y and x respectively represent the peak area and concentration. When estimating production per liter the concentration value was then extrapolated, considering the crude extract of 20 mL of media is concentrated to a final volume of 0.3 mL.

**Table S1.** Predicted spectra of dehydromicroperfuranone based on CFM-ID [4] is compared to LC-MS/MS ($MS^2$) spectra of m/z 265 at different fragmentation events.

<table>
<tr><td>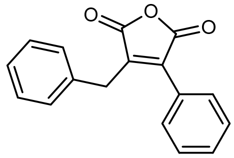<br>Dehydromicroperfuranone</td><td colspan="4">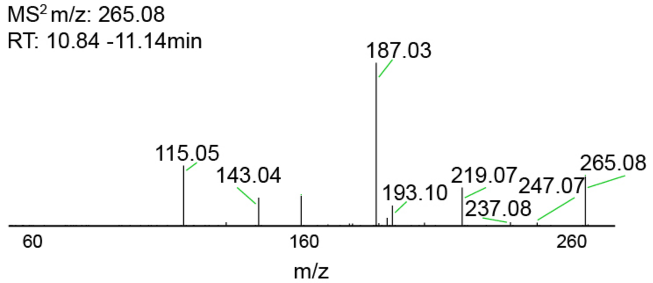
</td></tr>
<tr><td>Theoretical fragment m/z by CFM-ID</td><td>Observed m/z</td><td>Error (ppm)</td><td>Mass difference (Da)</td><td>Proposed structure by CFM-ID</td></tr>
<tr><td>265.0859207</td><td>265.079</td><td>-26.1080659</td><td>0.0069207</td><td>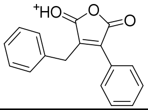</td></tr>
<tr><td>247.075356</td><td>247.0688</td><td>-26.53511896</td><td>0.006556</td><td>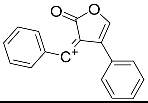</td></tr>
<tr><td>237.0910061</td><td>237.0847</td><td>-26.598511</td><td>0.0063061</td><td>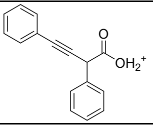</td></tr>
<tr><td>219.0804414</td><td>219.0745</td><td>-27.12045446</td><td>0.0059414</td><td>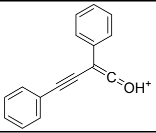</td></tr>
<tr><td>193.1011768</td><td>193.0959</td><td>-27.32735392</td><td>0.0052768</td><td>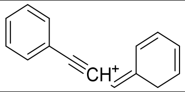</td></tr>
<tr><td>187.0389705</td><td>187.0337</td><td>-28.17941366</td><td>0.0052705</td><td>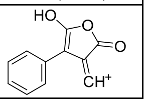</td></tr>
<tr><td>143.0491413</td><td>143.0451</td><td>-28.25192894</td><td>0.0040413</td><td>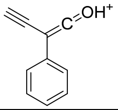</td></tr>
<tr><td>115.0542266</td><td>115.0509</td><td>-28.91415886</td><td>0.0033266</td><td>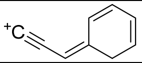</td></tr>
</table>

**Table S2.** $^{1}$H and $^{13}$C NMR data for dehydromicroperfuranone (**2**) purified as peaks **I-II** and peak **III** showing identical chemical shifts in chloroform-*d* (Figures S9–12). The previously reported NMR chemical shifts for **2** [5] are shown in the table for comparison. a: overlapping peaks, or tentatively assigned based on prediction.

| | **2** in Chloroform-*d* Ref. [5] | | **Peaks I-II** in Chloroform-*d* This work | | **Peak III** in Chloroform-*d* This work | |
|---|---|---|---|---|---|---|
| Carbon No. | $^{13}$C NMR | $^{1}$H NMR | $^{13}$C NMR | $^{1}$H NMR | $^{13}$C NMR | $^{1}$H NMR |
| 2 | 165.8 (s) | - | 166.0 (s) | - | 165.9 (s) | - |
| 3 | 141.0 (s) | - | 141.2 (s) | - | 141.3 (s) | - |
| 4 | 135.5 (s) | - | 135.5 (s) | | 135.6 (s) | - |
| 5 | 164.8 (s) | | 165.0 (d) | - | 165.0 (s) | - |
| 6 | 127.1 (s) | - | 127.2 (s) | - | 127.3 (s) | - |
| 7 | 128.4 (d) | 7.15 (5H, br s)[a] | 129.23 (d) | 7.21 (m)[a] | 129.3 (d)[a] | 7.21 (m)[a] |
| 8 | 129.0 (d) | 7.44 (5H, br s)[a] | 129.17 (d) | 7.63 (m)[a] | 129.2 (d)[a] | 7.63 (m)[a] |
| 9 | 130.2 (d) | 7.44 (5H, br s)[a] | 131.3 (d) | 7.55 (m)[a] | 131.4 (s) | 7.55 (m) |
| 10 | 129.0 (d) | 7.44 (5H, br s)[a] | 129.17 (d) | 7.63 (m)[a] | 129.2 (d)[a] | 7.63 (m)[a] |
| 11 | 128.4 (d) | 7.15 (5H, br s)[a] | 129.23 (d) | 7.21 (m)[a] | 129.3 (d)[a] | 7.21 (m)[a] |
| 12 | 30.4 (t) | 3.93 (2H, s) | 30.6 (s) | 4.03 (2H, s) | 30.6 (s) | 4.05 (2H, s) |
| 13 | 140.6 (s) | - | 140.8 (s) | - | 140.8 (s) | - |
| 14 | 129.3 (d) | 7.15 (5H, br s)[a] | 128.6 (d) | 7.34 (m)[a] | 128.6 (d) | 7.34 (m) |
| 15 | 129.0 (d) | 7.44 (5H, br s)[a] | 129.5 (d) | 7.53 (m)[a] | 129.5 (d)[a] | 7.53 (m)[a] |
| 16 | 127.3 (s) | 7.15 (5H, br s)[a] | 127.5 (d) | 7.29 (m)[a] | 127.5 (s) | 7.28 (m) |
| 17 | 129.0 (d) | 7.44 (5H, br s)[a] | 129.5 (d) | 7.53 (m)[a] | 129.5 (d)[a] | 7.53 (m)[a] |
| 18 | 129.3 (d) | 7.15 (5H, br s)[a] | 128.6 (d) | 7.34 (m)[a] | 128.6 (d) | 7.34 (m) |

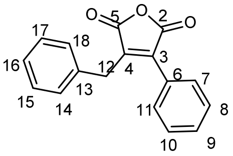


Dehydromicroperfuranone (2)

**Table S3.** $^{1}H$ and $^{13}C$ NMR data for microperfuranone (**1**) in chloroform-*d* (Figures S18–19). The previously reported NMR chemical shifts for **1** [6,7] are shown in the table for comparison. a: overlapping peaks, or tentatively assigned based on prediction.

| Carbon No. | **1** in acetone-$d_6$ Ref. 1 [6] | | **1** in Chloroform-*d* Ref. 2 [7] | | **1** in Chloroform-*d* This work | |
|---|---|---|---|---|---|---|
| | $^{13}C$ NMR | $^{1}H$ NMR | $^{13}C$ NMR | $^{1}H$ NMR | $^{13}C$ NMR | $^{1}H$ NMR |
| 2 | 170.9 (s) | - | 171.6 (s) | | 171.1 (s) | - |
| 3 | 130.8 (s) | - | 129.7 (s) | | 130.0 (s) | - |
| 4 | 159.6 (s) | - | 158.9 (s) | | 158.5 (s) | |
| 5 | 97.7 (d) | 5.98 (br s) | 97.2 (d) | 5.89 (br s) | 96.7 (d) | 5.97 (br s) |
| 5-OH | - | 6.90 (br s) | | N/A | - | N/A |
| 6 | 130.2 (s) | - | 129.1 (s) | | 129.1 (s) | - |
| 7 | 129.9 (d) | 7.54 (br d, 6.7) | 129.0 (s) | N/A | 129.2 (d) | 7.57 (m)[a] |
| 8 | 129.3 (d) | 7.46 (m) | 128.6 (d) | N/A | 128.8 (d) | 7.50 (m)[a] |
| 9 | 129.6 (d) | 7.43 (m) | 128.9 (d) | N/A | 128.9 (d) | 7.52 (m)[a] |
| 10 | 129.3 (d) | 7.46 (m) | 128.6 (d) | N/A | 128.8 (d) | 7.50 (m)[a] |
| 11 | 129.9 (d) | 7.54 (br d, 6.7) | 129.0 (s) | N/A | 129.2 (d) | 7.57 (m)[a] |
| 12 | 32.9 (t) | 3.97 (2H, br s) | 32.3 (s) | 3.91 (2H, br s) | 32.3 (s) | 3.97 (2H, br s) |
| 13 | 137.5 (s) | - | 136.0 (s) | | 136.1 (s) | - |
| 14 | 129.6 (d) | 7.30 (m) | 128.8 (d) | N/A | 128.9 (d) | 7.26 (m)[a] |
| 15 | 129.7 (d) | 7.26 (m) | 128.9 (d) | N/A | 129.1 (d) | 7.34 (m)[a] |
| 16 | 127.7 (d) | 7.22 (m) | 127.1 (d) | N/A | 127.3 (d) | 7.33 (m)[a] |
| 17 | 129.7 (d) | 7.26 (m) | 128.9 (d) | N/A | 129.1 (d) | 7.34 (m)[a] |
| 18 | 129.6 (d) | 7.30 (m) | 128.8 (d) | | 128.9 (d) | 7.26 (m)[a] |

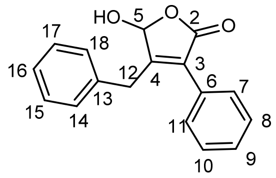


Microperfuranone (1)

**Note S1 : Sequence of crRNA/sgRNA expression cassettes cloning sites**

**> Cloning site of $P_{gpdA}$ *LbCas12a* crRNA expression cassette**

**(…)ATCTTCCCATCCAAGAACCTTTAATC**AAGCTTATCGATACCGTCGACCTCGACTCTAGAGGATCG***AATTTCTACTAAGTGTAGAT***GGAGACGAGAATTCCGTCTCC***AATTTCTACTAAGTGTAGAT***ATCTTCGAGGGGGGGCCCGGTACCGCCCCGTCCGGTCCTGCCCGTCACCGA**GATCCACTTAACGTTACTGAAATCAT(…)**

*A. nidulans* $P_{gpdA}$ TSS(bold underlined) and *gpdA* 5'UTR (bold green underlined). LbCas12a scaffold (blue italics). BsmbI sites (underlined).EcoRI site (red). **TrpC Terminator (black bold)**

**> Cloning site of $P_{U3}$ *SpCas9* sgRNA expression cassette**

*TTAATTAA***(...)CAAGTCAGAACATTTTGCTAACAGC**AGAGACGGGCGCCGCTACAGGGCGCGTCCCATTCGCCATTCAGGCTGCGCAACTGTTGGGAAGGGCGATCGGTGCGGGCCTCTTCGCTATTACGCCAGCTGGCGAAAGGGGGATGTGCTGCAAGGCGCGTCTCC*GTTTTAGAGCTAGAAATAGCAAGTTAAAATAAGGCTAGTCCGTTATCAACTTGAAAAAGTGGCACCGAGTCGGTGC***TTTTTTTT**CC*GCGGCCGC*CTGCAGGTCGACCATA(…)

PacI site (orange italics underlined) ***A. fumigatus U3* promoter (bold green)**. *SpCas9 sgRNA scaffold (blue italics).* BsmbI sites (underlined).LacZ fragment (red). Poly T terminator (bold). NotI site (purple italics underlined)
